# Export of discarded splicing intermediates requires mRNA export factors and the nuclear basket

**DOI:** 10.1101/2022.10.13.511998

**Authors:** Yi Zeng, Jonathan P. Staley

## Abstract

To promote fidelity in nuclear pre-mRNA splicing, the spliceosome rejects and discards suboptimal splicing substrates after they have engaged the spliceosome. Although nuclear quality control mechanisms have been proposed to retain immature mRNPs, evidence indicates that discarded splicing substrates, including lariat intermediates, do export to the cytoplasm, as indicated by their translation and degradation by cytoplasmic nucleases. However, the mechanism for exporting these species has remained unknown. By single molecule (sm) RNA FISH in budding yeast, we have directly observed the nuclear export of lariat intermediates. Further, by crosslinking, export reporter assays, and smRNA FISH, we have demonstrated that the export of lariat intermediates requires the general mRNA export receptor Mex67p and three of its mRNA export adapter proteins, Nab2p, Yra1p, and Nlp3, establishing that mRNAs and lariat intermediates share the same export machinery. Unexpectedly, the export of lariat intermediates, but not mRNA, requires an interaction between Nab2p and Mlp1p, a nuclear basket component implicated in retaining immature mRNPs, including unspliced pre-mRNA, in the nucleus of budding yeast. Finally, the export of lariat intermediates, like mRNA, relies on the E3 ubiquitin ligase Tom1p and its target sites in Yra1p. Overall, our data indicate that the nuclear basket can promote, rather than antagonize, the export of an immature mRNP. Further, our data imply that the export of discarded lariat intermediates requires both Mlp1p-dependent docking onto the nuclear basket and subsequent Tom1p-mediated undocking, a mechanism our data suggests functions in the export of mRNA also but in a manner obscured by redundant pathways.

## Introduction

In eukaryotes, the nuclear membrane uncouples translation from mRNA transcription and processing, thereby necessitating mRNA export through the nuclear pore complex (NPC; Petrovic et al., 2022), after transcription and RNA processing (Niño et al., 2013; Xie and Ren, 2019). Because the NPC establishes a permeability barrier, the export of such substrates requires the binding of export receptors to enable passage through the pore (Petrovic et al., 2022). The export of most mRNAs requires the heterodimeric export receptor – Mex67p-Mtr2p in yeast or NXF1-NXT1 in humans (Grüter et al., 1998; Katahira et al., 1999; Segref et al., 1997). The binding of Mex67p-Mtr2p to mRNA is mediated by export adapters, which bind mRNA in a manner that is generally coupled to transcription and RNA processing (Ben-Yishay and Shav-Tal, 2019; Stewart, 2019). Such coupling imparts specificity to Mex67p-Mtr2p binding. In a complementary manner, nuclear retention and decay have been invoked as a quality control mechanism that restricts the export of some mRNA transcripts, including immature or improperly processed messages (Palazzo and Lee, 2018). For example, the exosome, the PAXT complex, and PAXT complex components, especially ZFH3C1, have been implicated in retention and decay of such RNAs (Hilleren et al., 2001; Kwiatek et al., 2023; Lee et al., 2022; Ogami et al., 2017; Silla et al., 2018). In addition, the nuclear basket has been implicated in budding yeast and humans in the nuclear retention of incompletely spliced transcripts (Coyle et al., 2011; Galy et al., 2004; Palancade et al., 2005; Rajanala and Nandicoori, 2012); however, an absolute requirement for such a nuclear-basket quality control mechanism is challenged by a number of observations that also raise questions concerning the actual role of factors implicated in nuclear retention at the nuclear basket (Aksenova et al., 2020; Iino, 2017; Lee et al., 2020; Mayas et al., 2010; Palazzo and Lee, 2018; Sayani and Chanfreau, 2012). We have gained insight into these issues by investigating the proofreading of pre-mRNA splicing.

Pre-mRNA splicing is catalyzed by the spliceosome, a ribonucleoprotein machine that comprises over 80 conserved proteins and five small nuclear RNAs (snRNAs) (Wilkinson et al., 2020). The snRNA components play key roles in recognizing the substrate and catalyzing intron excision through two transesterification reactions; in the first reaction, branching, the 2’ hydroxyl of the branch site attacks the 5’ splice site, yielding a lariat intermediate and a free 5’ exon, which in the second reaction, exon ligation, attacks the 3’ splice site to yield mRNA and an excised intron. Proofreading in splicing functions to enhance the specificity of splice site choice (Semlow and Staley, 2012). Grossly suboptimal splice sites never bind the spliceosome, but suboptimal splice sites do bind, necessitating downstream mechanisms to distinguish and reject such splice sites. Indeed, proofreading mechanisms have been implicated throughout the splicing cycle, including the phases of spliceosome assembly, activation, and catalysis. ATPases of the DEAD and DExH families of the helicase II superfamily play a central role in proofreading (De Bortoli et al., 2021; Semlow and Staley, 2012). In addition to driving the splicing cycle forward in the case of optimal splice sites, these ATPases also reject and discard splicing substrates having suboptimal splice sites. For example, the DExH ATPase Prp22p, in the case of an optimal substrate, promotes the release of mRNA from the spliceosome after exon ligation, but in the case of a suboptimal 3’ splice site, Prp22p rejects the splice site by undocking the splice site from the catalytic core before exon ligation (Mayas et al., 2006; Semlow et al., 2016). If an alternative, optimal 3’ splice site does not engage the spliceosome, then a second DExH ATPase Prp43p, which acts as a general terminator of splicing, disassembles the spliceosome, discarding the suboptimal substrate at the intermediate stage (Mayas et al., 2010; Pandit et al., 2006). As a consequence of such proofreading pathways, any mechanism proposed to enforce robust quality control of mRNA export would need to distinguish against not only pre-mRNA that failed to engage the spliceosome but also pre-mRNA that did engage the spliceosome but subsequently suffered discard at either the pre-mRNA or the intermediate stage. A robust quality control mechanism to discriminate against the nuclear export of such species has been thought to be essential to preclude the translation of such species into truncated protein products (Coyle et al., 2011; Galy et al., 2004; Rajanala and Nandicoori, 2012) (but see below).

Specific factors have been implicated in coupling transcription and RNA processing to mRNA export and in the nuclear retention of immature species to establish specificity in mRNA export (Tutucci and Stutz, 2011; Schmid and Jensen, 2018; Palazzo and Lee, 2018). For example, the TREX complex, which functions in transcription elongation and mRNA export, favors the export of spliced RNA (Masuda et al., 2005; Tutucci and Stutz, 2011). Further, reporter assays have implicated Mlp1p (TPR in humans), a subunit of the nuclear basket, as a quality control factor for retaining faulty RNA transcripts, especially unspliced pre-mRNAs, in the nucleus (Coyle et al., 2011; Galy et al., 2004; Palancade et al., 2005; Rajanala and Nandicoori, 2012). However, several observations question the requirement for strict quality control mechanisms in mRNA export. First, several studies found that Mlp1p/TPR does not act as a general quality control factor in broadly retaining pre-mRNA in the nucleus and instead promotes mRNA export in some cases (Bangs et al., 1998; Green et al., 2003; Bae et al., 2009; Xu et al., 2007; Lee et al., 2020; Zuckerman et al., 2020; Aksenova et al., 2020; Iino, 2017; Sayani and Chanfreau, 2012). Further, evidence indicates that pre-mRNA and splicing intermediates can be localized to the cytoplasm in both budding yeast and humans (Mayas et al., 2010; Harigaya and Parker, 2012; Sayani and Chanfreau, 2012; Carvalho et al., 2017; Talhouarne and Gall, 2018; Hilleren and Parker, 2003; Legrain and Rosbash, 1989). Lastly, disabling the nonsense-mediated decay machinery stabilizes pre-mRNAs in the cytoplasm (Sayani and Chanfreau, 2012). These observations raise questions about a strict requirement for quality control in mRNA export and the role of factors implicated in nuclear retention, in particular, the nuclear basket. Further, the cytoplasmic localization of splicing intermediates raises questions concerning the pathway utilized for export and the relation of this pathway to mRNA export, because, for example, a cleaved 5’ exon lacks a poly(A) tail, and a lariat intermediate lacks a cap, both features of mRNA implicated in promoting export.

In yeast, several mRNA export adapters have been identified, including the SR-like protein Npl3p, Nab2p, the TREX-2 complex, and the TREX complex, featuring Yra1p (Tutucci and Stutz, 2011). In addition to interacting with mRNA and Mex67p-Mtr2p, these adapters often interact with additional factors. For instance, the N-terminal domain of Nab2, a poly-A tail binding protein (Anderson et al., 1993), interacts directly with the C-terminal domain of Mlp1p (Grant et al., 2008). Yra1p has also been implicated in binding to the nuclear basket, through interactions with Mlp1p and Mlp2p in an RNA-dependent manner (Vinciguerra et al., 2005). Interestingly, the E3 ubiquitin ligase Tom1p ubiquitylates Yra1p and then dissociates Yra1p from mRNAs in the nucleus (Iglesias et al., 2010). Deletion of *TOM1* inhibits mRNA export at high temperature (Duncan et al., 2000), and genetic assays imply that deletion of *TOM1* leads to nuclear retention of mRNA in an Mlp1p and Mlp2p dependent manner (Iglesias et al., 2010).

Mlp1p and Mlp2p compose the elongated fiber-like structures of the nuclear basket (Strambio-de-Castillia et al., 1999; Niepel et al., 2013; Kim et al., 2018). The N-termini of these proteins are anchored to the nuclear pore, and their C-termini extend roughly 300 nm into the nuclear interior (Bangs et al., 1998; Kim et al., 2018). In addition to the purported roles in retaining immature RNAs in the nucleus, Mlp1p and TPR have been implicated in mRNA export in *S. pombe*, *Arabidopsis*, and mammalian cells as perturbations of these proteins result in the accumulation of poly(A)+ RNA in the nucleus and/or defects in export of specific mRNAs to the cytoplasm (Aksenova et al., 2020; Bae et al., 2009; Bhat et al., 2023; Lee et al., 2020; Li et al., 2021; Shibata et al., 2002; Umlauf et al., 2013; Xu et al., 2007; Zuckerman et al., 2020). Indeed, rapid depletion of TPR and NXF1 results in similar mRNA export defects (Aksenova et al., 2020; Zuckerman et al., 2020), supporting an mRNA export role for TPR, which microscopy implies functions in mRNA docking to the NPC (Aksenova et al., 2020; Lee et al., 2020; Li et al., 2021). Consistent with a role in mRNA export, Mlp1p/2p/TPR interact not only with the mRNA export adapters Nab2p and Yra1p but also Npl3p and the TREX-2 complex (Ashkenazy-Titelman et al., 2020; Fasken et al., 2008; Green et al., 2003; Vinciguerra et al., 2005). However, Mlp1p/TPR may not play a general role in mRNA export, because in budding yeast *MLP1* is not essential for growth or mRNA export (Bangs et al., 1998; Green et al., 2003; Kosova et al., 2000), and in mammals TPR is only required to export a subset of poly(A)+ mRNAs (Umlauf et al., 2013) that are short, intron poor, or GC poor (Lee et al., 2020; Zuckerman et al., 2020). The lack of a general role for Mlp1p and Mlp2p in mRNA export has suggested a broader role for Mlp1p and its orthologs in nuclear retention of immature substrates, but the role of Mlp1p in both export and retention remains under investigation.

In this work, we have gained a unique perspective into the role of Mlp1p in mRNA export and quality control through an investigation of the export pathway for discarded lariat intermediates, in budding yeast where export studies are not complicated by open mitosis. We found that, like the export of mRNA, the export of discarded, lariat intermediates require the export receptor Mex67p and three of its adapters, Yra1p, Nab2p, and Npl3p. Unlike mRNA export, lariat intermediate export requires Mlp1p and an interaction between Mlp1p and Nab2p. Further, lariat intermediate export required, even at permissive temperature, Tom1p-mediated ubiquitylation of Yra1p. Unexpectedly, our data argue against a general role for Mlp1p in quality control through nuclear retention of immature RNAs and instead highlight a role for Mlp1p in exporting immature RNAs. Specifically, our data suggest a model in which the export of lariat intermediates first requires docking onto the nuclear basket and then undocking to permit transit through the nuclear pore, a pathway that likely also operates in the case of mRNA export but in a manner that is normally masked by redundant pathways.

## Results

### Export of lariat intermediates requires the mRNA export factor Mex67p

To investigate the mechanism of lariat intermediate export in *Saccharomyces cerevisiae*, we tested whether the canonical mRNA export factor Mex67p is required, given that the association of its adapters on a transcript initiates co-transcriptionally. First, to observe the subcellular localization of lariat intermediates, we utilized a *lacZ*-expressing *ACT1* export reporter that includes a branch site (br) A-to-G mutation and thus accumulates lariat intermediates that localize to the cytoplasm, based on indirect assays (**Fig. 1A, B**; Mayas et al., 2010). To define the subcellular localization of reporters directly, we targeted the *lacZ* gene in the 3’ exon of the derived reporter, named brG, and the wild-type control, named brA, for smRNA FISH. To resolve individual foci and thereby allow accurate, subcellular RNA counting, we modified the reporter by replacing its strong promoter with the weak *pSTE5* promoter. Consistent with previous indirect assays (Mayas et al., 2010), in a wild-type strain at 30 °C the vast majority of the brG reporter localized to the cytoplasm, as for the brA reporter, with only a minor fraction in the nucleus (17% ±1% (SEM) and 17% ±1%, respectively; **Fig. 1C**). By primer extension analysis, the brG mutation increases lariat intermediate levels by 7-fold (**Fig. 1B**); although the mutation also increases pre-mRNA levels by 2-fold, the pre-mRNA represents only 33% of all brG species (**Fig. 1B**), which is insufficient to account for the magnitude of the cytoplasmic signal from the brG reporter (83%), and a different reporter that accumulates lariat intermediate exclusively also localizes to the cytoplasm by smRNA FISH (see below). These data provide direct evidence for the nuclear export of lariat intermediates.

**Figure 1.**
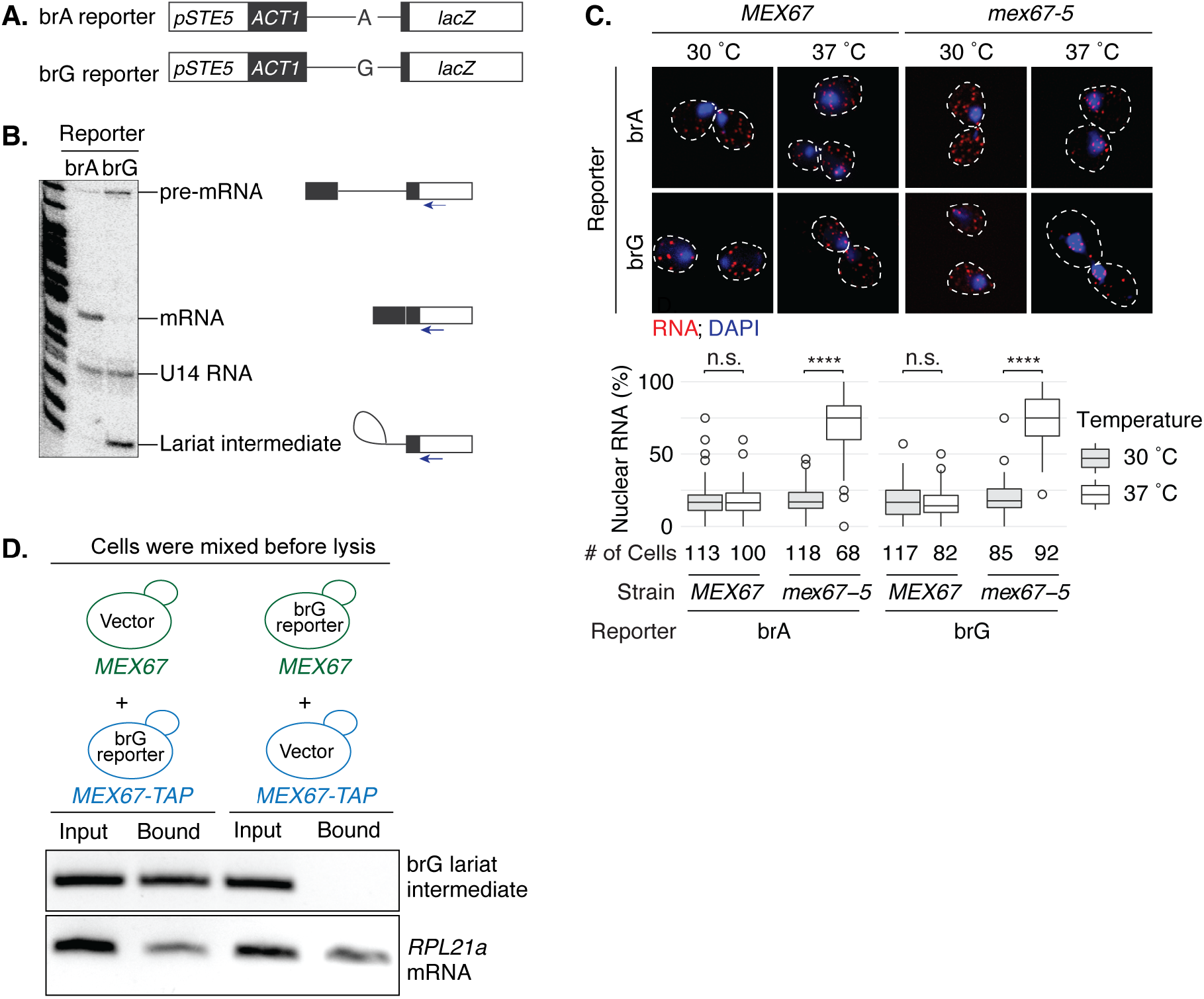
Export of a lariat intermediate requires the mRNA export factor Mex67p. **A.** Schematic representation of the brA and brG export reporters. **B.** The brG reporter accumulates lariat intermediate, as well as pre-mRNA to a lesser extent, as shown by extension of a primer that anneals to the downstream exon. The migration of the indicated splicing species is shown. **C.** By smRNA FISH, the export of the brG reporter is impeded by the *mex67-5* mutant at the non-permissive temperature. Wild-type or *mex67-5* mutant cells transformed with the indicated reporters were shifted from 30 °C to 37 °C for 30 minutes. The fraction of nuclear RNA for each cell was quantitated and displayed as a box plot. The number of cells used for quantitation is indicated beneath each box plot. The *lacZ* region of the reporter transcript was targeted by Cy3-labeled probes; DNA was probed by DAPI. Representative cells are shown; dashed lines mark the cell boundary. **D.** By RNA co-IP, Mex67p interacts with lariat intermediates *in vivo*. Top, the indicated cells were mixed before lysis and co-IP. Bottom, the brG lariat intermediate or *RPL21a* mRNA was detected by RT-PCR before (Input) and after (Bound) IP of Mex67p-TAP, expressed from the indicated strains (*MEX67-TAP*). The *p*-values were calculated by Mann-Whitney test; n.s. (not significant), *p* > 0.05; *, *p* ≤ 0.05; **, *p* ≤ 0.01; **\*\*\***, *p* ≤ 0.001; ****, *p* ≤ 0.0001.

Next, we used smRNA FISH and RNA FISH to examine whether the export of lariat intermediates required Mex67p. Indeed, in a temperature-sensitive *mex67-5* mutant that blocks mRNA export at the non-permissive temperature of 37 °C (Segref et al., 1997), both the brG and brA reporters accumulated in the nucleus after a temperature shift to 37 °C (**Fig. 1C; Fig. S1A, B**). Specifically, by smRNA FISH, the localization of the brG reporter in the nuclei of *mex67-5* cells increased 3.9-fold from 19% ±1% to 74% ±2%, paralleling a 4.0-fold increase in brA-derived mRNAs from 18% ±1% to 71% ±3% (**Fig. 1C**); by RNA FISH, the localization of the brG reporter, like poly(A)+ RNA, increased similarly in nuclei (**Fig. S1B**). In contrast, a shift of wild-type *MEX67* cells did not significantly increase localization of the brG reporter in the nucleus (16% ±1% at 37 °C compared to 17% ±1% at 30 °C), as for brA-derived mRNA and poly(A)+ RNA (**Fig. 1C; Fig. S1B**). In contrast with the *mex67-5* mutant, the tRNA export mutant *los1Δ* did not affect the cytoplasmic localization of the brG reporter, indicating that the export of lariat intermediates is not promiscuous and requires a specific export pathway (**Fig. S1C**). Together, our results indicate that the export of lariat intermediates requires the general mRNA export factor Mex67p.

To test whether Mex67p interacts with lariat intermediates directly, we assayed for the interaction of Mex67p with lariat intermediates *in vivo*. We performed RNA co-immunoprecipitation (co-IP) from extracts of *MEX67-GFP* cells expressing the brG reporter and found that Mex67p-GFP did co-immunoprecipitate lariat intermediates (**Fig. S1D**). To rule out that the interaction formed *in vitro*, we pre-mixed untagged *MEX67* cells and TAP-tagged *MEX67* cells, lysed the cell mixture, and then performed RNA co-IP. When we expressed the reporter in the tagged cells, we again observed co-immunoprecipitation of lariat intermediates with tagged *MEX67*, but when we expressed the brG reporter in the untagged cells, we did not observe co-immunoprecipitation (**Fig. 1D**); by contrast, endogenous *RPL21a* mRNA co-immunoprecipitated with tagged *MEX67* in either condition. These data verify that Mex67p interacts with lariat intermediates *in vivo* and support a direct role for Mex67p in lariat intermediate export.

### Export of lariat intermediates requires Mex67p export adaptors

In budding yeast, Mex67p is recruited to mRNA transcripts by three different export adaptors, Yra1p, Nab2p, and/or Npl3p (Gilbert and Guthrie, 2004; Hurt et al., 2004; Iglesias et al., 2010; Zenklusen et al., 2001). Therefore, we tested whether any of these adaptors is required for the export of lariat intermediates. First, to test for the requirement of Yra1p in the export of lariat intermediates, we tested whether the temperature-sensitive *GFP-yra1-8* mutant, which blocks mRNA export at 37 °C (Zenklusen et al., 2002), disrupts the export of lariat intermediates at 37 °C. Specifically, we transformed the brG and brA reporters into *GFP-yra1-8* cells, shifted the cells to 37 °C, and then assessed the cellular localization of the reporters by smRNA FISH. As for the nuclear levels of the brA reporter, the nuclear levels of the brG reporter did not increase in the control, wild-type *YRA1* cells, but they did increase 2.6-fold in the *GFP-yra1-8* cells, from 28% ±1% to 72% ±4% (**Fig. 2A, B**), thereby establishing evidence that the export of lariat intermediates requires Yra1p. To test whether lariat intermediates interact directly with Yra1p, we expressed the brG reporter in *HA-YRA1* cells, formaldehyde crosslinked the cells, performed denaturing RNA co-IP from extracts, and then assayed for RNA by RT-qPCR using primers specifically targeting lariat intermediates. Indeed, HA-Yra1p not only co-immunoprecipitated endogenous *RPL21a* mRNA, as expected, but also brG-derived lariat intermediates (**Fig. 2C**). These data support a direct role for Yra1p in the export of lariat intermediates.

**Figure 2.**
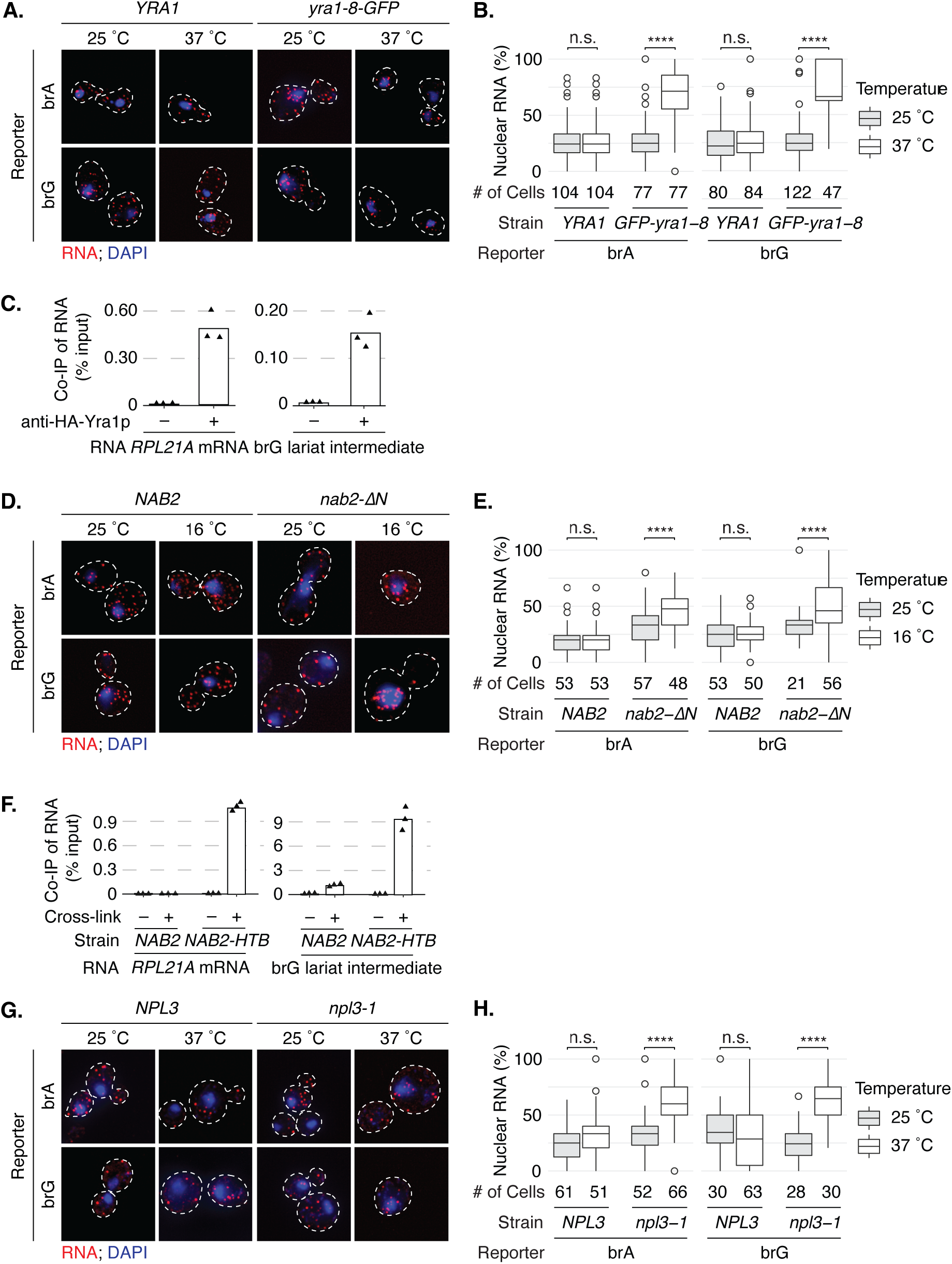
Export of a lariat intermediate requires the mRNA export adapters Yra1p, Nab2p, and Npl3p. **A, B.** By smRNA FISH, export of the brG reporter requires *YRA1*. Wild-type or *yra1-8-GFP* mutant cells were shifted from 25 °C to 37 °C for 2 hours. Panel **A** shows representative images. Panel **B** shows the quantitation of the nuclear fraction of reporter RNA. **C.** By RNA co-IP, Yra1p interacts directly with lariat intermediates. After formaldehyde crosslinking, HA-Yra1p was immunoprecipitated from cell extracts under denaturing conditions with or without antibodies, and then associated brG lariat intermediate or *RPL21a* mRNA was detected by RT-qPCR. The levels of immunoprecipitated RNA are shown as a percentage of input for three technical replicates; the height of the bar indicates the mean. **D, E.** By smRNA FISH, the export of the brG reporter requires *NAB2*. Wild-type or *nab2-ΔN* mutant cells were shifted from 30 °C to 16 °C for 2 hours. Panel **D** shows representative images; panel **E** shows quantitation of the nuclear fraction of reporter RNA. **F.** By RNA co-IP, Nab2p interacts directly with lariat intermediates. HTB-tagged *NAB2-HTB* cells and untagged control cells were treated with and without UV, Nab2p-HTB was pulled down from cell extracts under denaturing conditions using Ni-NTA agarose, and then associated brG lariat intermediate or *RPL21a* mRNA was assayed by qRT-PCR. The levels of pulled-down RNA were quantitated and illustrated as in panel **C**. **G, H.** By smRNA FISH, the export of the brG reporter requires *NPL3*. Wild-type or *npl3-1* mutant cells were shifted from 25 °C to 37 °C for 2 hours. Panel **G** shows representative images; panel **H** shows the quantitation of the nuclear fraction of reporter RNA. Throughout, for smRNA FISH cells were probed and demarked as in Fig. 1C (top panel); the nuclear fraction was calculated and displayed as in Fig. 1C (bottom panel).

Next, to test for a requirement of Nab2p in the export of lariat intermediates, we assessed by smRNA FISH whether the export of the brG reporter was compromised in a cold-sensitive *nab2-ΔN* mutant, which accumulates poly(A)+ RNA in the nucleus at 16 °C (Marfatia et al., 2003). Indeed, when *nab2-ΔN* cells expressing the reporters were shifted to 16 °C, the nuclear levels of the brG reporter increased 1.5-fold from 35% ±4% to 52% ±3%, similar to the increase of 1.5-fold from 32% ±2% to 47% ±2% for the brA reporter (**Fig. 2D, E**), suggesting that Nab2p is also required for the export of lariat intermediates. To test whether Nab2p interacts directly with lariat intermediates, we expressed the brG reporter in HTB-tagged *NAB2-HTB* cells, UV-crosslinked the cells, performed denaturing RNA pull-down from extracts, and then assayed for associated RNAs by RT-qPCR. Indeed, Nab2p-HTB not only pulled down *RPL21a* mRNA in a UV-dependent manner, as expected, but also brG-derived lariat intermediate (**Fig. 2F**). These data support a direct role for Nab2p in the export of lariat intermediates.

Finally, to examine whether Npl3p plays a role in the export of lariat intermediates, we tested whether a temperature-sensitive *npl3-1* mutant, which displays a strong defect in mRNA export after a shift to 37 °C (Lee et al., 1996), blocks the export of lariat intermediates. After a shift from 25 °C to 37 °C for 2 hours, the nuclear localization of the brG reporter in the *npl3-1* cells increased 2.6-fold from 24% ±3% to 63% ±4%, similar to the increase of 1.8-fold from 33% ±2% to 60% ±3% for the brA reporter (**Fig. 2G, H**). Unfortunately, an Npl3-TAP co-IP was unsuccessful, so we were unable to confirm direct binding to the brG reporter. Nevertheless, these data suggest that, like Yra1p and Nab2p, Npl3p contributes to the export of lariat intermediates. Taken together, these results establish evidence that the export of lariat intermediates requires Mex67p and its adaptors and therefore the canonical mRNA export pathway.

### Efficient export of lariat intermediates requires the nuclear basket component Mlp1p

Because previous studies have determined that Nab2p physically interacts with the nuclear basket factor Mlp1p and that the *nab2-ΔN* mutant lacks the Mlp1p-interacting domain (Marfatia et al., 2003; Grant et al., 2008), we hypothesized that Nab2p mediates lariat intermediate export through its interaction with Mlp1p, even though Mlp1p has been implicated as a quality-control factor that retains immature RNPs in the nucleus (Bonnet and Palancade, 2015; Galy et al., 2004; Vinciguerra et al., 2005). As a first test of this hypothesis, we assayed whether the export of the brG reporter requires Mlp1p. To assay export efficiency, we used our previously described *lacZ*-based export reporter having an internal ribosomal entry site (IRES) in the second exon just upstream of *lacZ*, which enables the translation of exported species, whether a pre-mRNA, lariat intermediate, or mRNA species (**Fig. 3A**; Mayas et al., 2010). Indeed, deletion of *MLP1* significantly reduced the β-galactosidase activity of the brG-IRES reporter by 50%; by contrast deletion of *MLP1* did not significantly affect the β-galactosidase activity of the brA-IRES reporter (**Fig. 3B**). This reduced β-galactosidase activity in *mlp1Δ* cells cannot be accounted for by changes in RNA levels, which were not perturbed (**Fig. S2B, C**). Similarly, in *mlp1Δ* cells a G1a reporter that accumulates lariat intermediates showed reduced β-galactosidase activity (**Fig. S2B, C, D**). These data are consistent with a requirement for Mlp1p in the export of lariat intermediates (see below).

**Figure 3.**
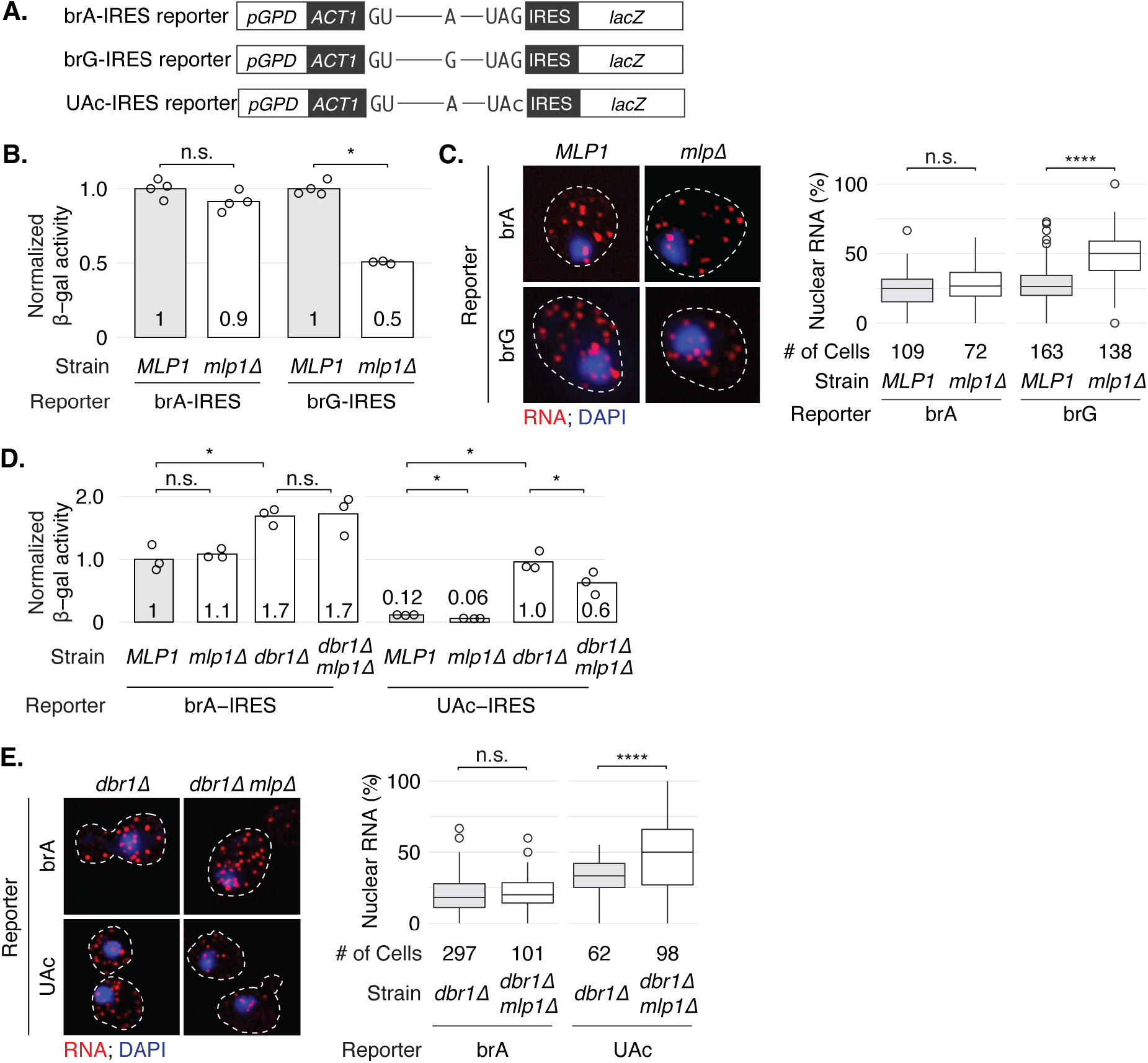
The export of lariat intermediates requires Mlp1p. **A.** Schematic representation of the *ACT1-IRES-lacZ* export reporters. **B, C.** Efficient export of the brG but not the brA reporter requires *MLP1*. In panel **B**, *MLP1* or *mlp1Δ* cells were grown at 30 °C, and the cytoplasmic localization of the reporters was assayed by measuring the β-galactosidase activity of cell extracts. The normalized β-galactosidase activities of four technical replicates are shown; the height of the bar and the number indicate the mean activity, normalized to the mean activity of the same reporter in the *MLP1* strain. In panel **C**, *MLP1* or *mlp1Δ* cells were grown at 30 °C, and the cytoplasmic localization of the reporters was assayed directly by smRNA FISH (left); the nuclear fraction of reporter RNA was quantitated (right). **D, E**. The export of the UAc lariat intermediate is comprised by the *mlp1Δ* mutation. In panel **D**, *MLP1*, *mlp1Δ*, *dbr1Δ*, or *mlp1Δ dbr1Δ* cells were grown at 30 °C, and the cytoplasmic localization of the brA- and UAc-IRES reporters was assayed by measuring β-galactosidase activity of cell extracts. The β-galactosidase activity was quantitated as in B; values were normalized to the brA-IRES reporter in the wild-type strain (*MLP1*). In panel **E**, mutant *dbr1Δ* or double mutant *mlp1Δ dbr1Δ* cells were grown at 30 °C and then assayed for reporter localization by smRNA FISH (left); the nuclear fraction of reporter RNA was quantitated (right). Throughout, for smRNA FISH, cells were probed and demarked as in Fig. 1C; the nuclear fraction was calculated and displayed as in Fig. 1C. The *p*-values were calculated by Mann-Whitney test and represented by asterisks as in Fig. 1.

To determine directly whether *MLP1* promotes the export of the brG reporter, we used smRNA FISH to examine the cellular localization of the reporter in *mlp1Δ* cells. Consistent with our findings by the β-galactosidase assay, the *mlp1Δ* mutation decreased the cytoplasmic signal and concomitantly increased the nuclear signal of the brG reporter; the nuclear fraction increased by 1.8-fold – from 27% ±1% in wild-type *MLP1* cells to 50% ±2% in *mlp1Δ* cells (**Fig. 3C**). By contrast, the *mlp1Δ* mutation did not impact the subcellular localization of the brA reporter (**Fig. 3C**). These data verify directly that *MLP1* promotes the export of the brG reporter, and the data are consistent with a role for Mlp1p in the export of lariat intermediates.

Because the brG reporter accumulates some pre-mRNA (38%), in addition to lariat intermediate (58%; **Fig. S2B, C**) and the *mlp1Δ* mutant only partially compromised export of the brG reporter (by 50%; **Fig. 3B**), we tested whether the dependence of the brG reporter on Mlp1p might reflect a dependence of the export pre-mRNA export, instead of lariat intermediate, on Mlp1p. To test this possibility, we assessed the impact of the *mlp1Δ* mutation on the export of an isogenic reporter having a G1c mutation at the 5’ splice site, a mutation that accumulates similar levels of pre-mRNA (2.4-fold increased, relative to wild type; **Fig. S2B, C**) but no lariat intermediate; in fact, the levels of lariat intermediate are even lower than for a wild-type reporter (**Fig. S2B**). Significantly, the *mlp1Δ* mutation did not reduce the levels of β-galactosidase expressed from the G1c reporter, relative to the wild-type strain (**Fig. S2E)**. Similarly, in *mlp1Δ* cells, a brC reporter that similarly accumulates pre-mRNA but not lariat intermediate did not reduce β-galactosidase activity (**Fig. S2B, C, E**). These data provide strong evidence that the export of brG pre-mRNA is not dependent on Mlp1p; these data, therefore, imply that the efficient export of brG lariat intermediates is dependent on Mlp1p.

To test for a requirement for Mlp1p in the export of lariat intermediates explicitly, we assessed the impact of the *mlp1Δ* mutation on the export of an isogenic reporter having a UAc mutation at the 3’ splice site. Like the brG mutation, the UAc mutation compromises exon ligation; unlike the brG mutation, the UAc mutation does not substantially compromise the conversion of pre-mRNA to lariat intermediate, so UAc pre-mRNA levels are insignificant (e.g., **Fig. S3A**). However, whereas the brG lariat intermediate, having a non-consensus branch linkage, is resistant to debranching by Dbr1 and to subsequent turnover, the UAc lariat intermediate, with a consensus branch linkage, is subject to rapid debranching and turnover. Nevertheless, in a *dbr1Δ* strain the UAc lariat intermediate is stabilized (e.g., 4-fold in **Fig. S3A, B**), and this stabilized lariat intermediate is efficiently translated in the cytoplasm, as we have shown previously (**Fig. 3D**; Mayas et al., 2010). Thus, we tested in a *dbr1Δ* background whether the export of the UAc lariat intermediate requires *MLP1* for export. Indeed, the *mlp1Δ* mutation compromised export; specifically, whereas the *mlp1Δ* mutation did not significantly reduce the β-galactosidase activity of the brA-IRES reporter, the *mlp1Δ* mutation reduced the β-galactosidase activity of the UAc-IRES reporter by roughly 40% (**Fig. 3D**), similar to the brG reporter (**Fig. 3B**). Even in the wild-type *DBR1* strain, where low levels of UAc-IRES lariat intermediate accumulates relative to the brA-IRES control (**Fig. S3A, B)**, we detected a similar decrease in β-galactosidase activity for the UAc-but not the brA-IRES reporter (**Fig. 3D**). These data provide compelling evidence that *MLP1* is required explicitly and specifically for the efficient export of lariat intermediates.

To test directly for a requirement for *MLP1* in the export of the UAc lariat intermediates, we assayed for localization of the UAc reporter in the *dbr1Δ* background by smRNA FISH. Whereas the *mlp1Δ* mutation did not impact the subcellular localization of the brA reporter, the *mlp1Δ* mutation did shift the localization of the UAc reporter from the cytoplasm to the nucleus; specifically, we observed a shift in the nuclear fraction from 33% ±2% to 45% ±3% (**Fig. 3E**), similar to the shift of the brG reporter in a *DBR1* background (**Fig. 3C**); by contrast, the *mlp1Δ* mutation did not shift the localization of the brA reporter from the cytoplasm to the nucleus (**Fig. 3E**). These data provide direct evidence for a specific requirement for *MLP1* in efficient lariat intermediate export.

### Export of lariat intermediates requires an interaction between Nab2p and Mlp1p

As a further test of our hypothesis that Nab2p mediates lariat intermediate export through its interaction with Mlp1p, we probed more deeply for a requirement for the Nab2p-Mlp1p interface in the export of lariat intermediates. First, we tested for a requirement for the Mlp1p component of the interface utilizing the *mlp1-Δ1586-1768* mutation, which lacks the Nab2p-interacting domain. This mutation compromised the β-galactosidase activity of the brG-IRES reporter; specifically, whereas expression of wild-type *MLP1* in *mlp1Δ* cells fully rescued the β-galactosidase activity of the brG-IRES reporter, relative to *mlp1Δ* cells with a vector control, expression of *mlp1-Δ1586-1768* failed to rescue (**Fig. 4A**). By fluorescence microscopy, we confirmed that the mutated Mlp1p-Δ1586-1768 protein is localized properly to the nuclear periphery, as observed for Mlp1p (**Fig. S4A**; Galy et al., 2004; Niepel et al., 2013). These results demonstrate that efficient lariat intermediate export requires the interaction between Nab2p and Mlp1p.

**Figure 4.**
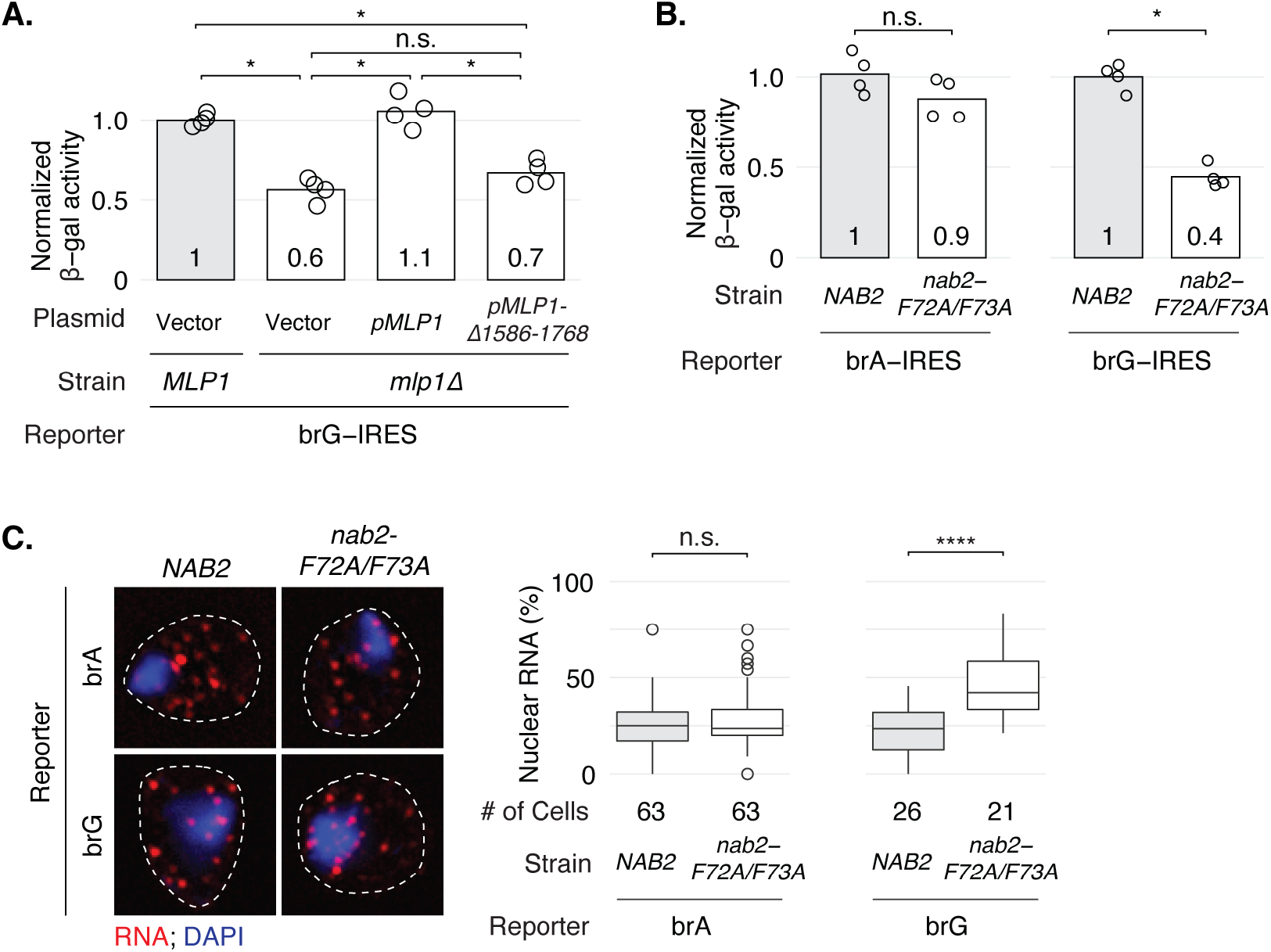
The export of lariat intermediates requires an interaction between Nab2p and Mlp1p. **A**. A deletion in the Nab2p-interacting domain of Mlp1p reduces the export of the brG reporter. Mutant *mlp1Δ* cells were transformed with a vector control, *MLP1*, or *mlp1Δ1586-1768*, grown at 30 °C, and then lysed to assay β-galactosidase activity of the brG reporter. The activity was quantitated as in Fig. 3B; values were normalized to the brG reporter in the *MLP1* strain transformed with a vector control. Panels **B** and **C** show that a double mutation in the Mlp1p-interacting domain of Nab2p reduces the export of the brG reporter, specifically. In panel **B**, *NAB2* or *nab2-F72AF73A* cells were grown at 30 °C and then lysed to assay β-galactosidase activity. The activity was quantitated as in Fig. 3B; values for each reporter were normalized to the mean activity of the same reporter in the *NAB2* strain. In panel **C,** *NAB2* or *nab2-F72AF73A* cells were grown at 30 °C and then assayed for reporter localization by smRNA FISH (left); cells were probed and demarked as in Fig. 1C. The nuclear fraction of reporter RNA was quantitated and displayed, on the right, as in Fig. 1C.

We showed above that the cold-sensitive *nab2-ΔN* mutant, which lacks the Mlp1p-interacting domain, compromises the export of the brG reporter at a low temperature (**Fig. 2D, E**), but this mutant also compromised the export of the brA reporter (**Fig. 2D, E**), in addition to poly(A)+ RNA (Grant et al., 2008) – phenotypes that do not parallel the phenotypes of *mlp1* mutants, so we next assayed whether lariat intermediate export was sensitive to a subtler *NAB2* double mutant, *nab2-F72A/F73A*, that also disrupts the Nab2p-Mlp1p interaction (Fasken et al., 2008; Grant et al., 2008) but displays no growth defect at any temperature (**Fig. S4B**). Indeed, although the *nab2-F72A/F73A* mutation did not significantly affect the β-galactosidase activity of the brA-IRES reporter, the *nab2-F72A/F73A* mutation did reduce the β-galactosidase activity of the brG-IRES reporter by 60% (**Fig. 4B**), quantitatively similar to the reduction of β-galactosidase activity observed for *mlp1Δ* or *mlp1-Δ1586-1768* mutant cells (**Fig. 3B**; **Fig. 4A**). The diminished β-galactosidase activity of the brG *lacZ* reporter in *nab2-F72A/F73A* cells did not result from changes in RNA levels, because the *nab2-F72A/F73A* mutant did not alter RNA levels of either reporter (**Fig. S4C, D**). These data are also consistent with the hypothesis that efficient lariat intermediate export requires the interaction between Nab2p and Mlp1p.

To determine directly whether the Nab2p-Mlp1p interaction promotes the export of lariat intermediates, we used smRNA FISH to examine the cellular localization of the brG reporter in *nab2-F72A/F73A* mutant cells. As in *mlp1Δ* cells (**Fig. 3C**), in *nab2-F72A/F73A* cells the nuclear fraction of the brG reporter increased by 2.1-fold, from 23% ±3% in wild-type *NAB2* cells to 48% ±4% in *nab2-F72A/F73A* cells, whereas the nuclear fraction of the brA reporter did not increase significantly (**Fig. 4C**). Together, these results establish that the export of lariat intermediates, in particular, requires the interaction between Nab2p and Mlp1p.

### Export of lariat intermediates requires Tom1p-mediated ubiquitylation of Yra1p

Yra1p is ubiquitylated by the E3 ligase Tom1p, and this ubiquitylation is required for efficient poly(A)+ mRNA export at a higher temperature (37 °C; Iglesias et al., 2010; Duncan et al., 2000). Additionally, based on genetic interactions of Tom1p or Yra1p with Mlp1p or Mlp2p, Tom1p has been proposed to surveil mRNAs for maturity at the nuclear basket (Iglesias et al., 2010). Thus, we investigated the impact of Tom1p on lariat intermediate export. First, we assayed for the consequences of deleting *TOM1* on the export of the brA- and brG-IRES reporters. At the permissive temperature (30 °C), whereas the *tom1Δ* mutation did not significantly reduce the β-galactosidase activity of the brA-IRES reporter, the deletion mutation did significantly reduce the activity of the brG-IRES reporter (**Fig. 5A**), suggesting unexpectedly that *TOM1, like MLP1,* promotes, rather than antagonizes, the export of incompletely processed mRNPs; furthermore, our data imply that Tom1p, like Mlp1p and its interaction with Nab2p (**Fig. 3**; **Fig. 4**), is particularly important for the export of incompletely processed mRNPs.

**Figure 5.**
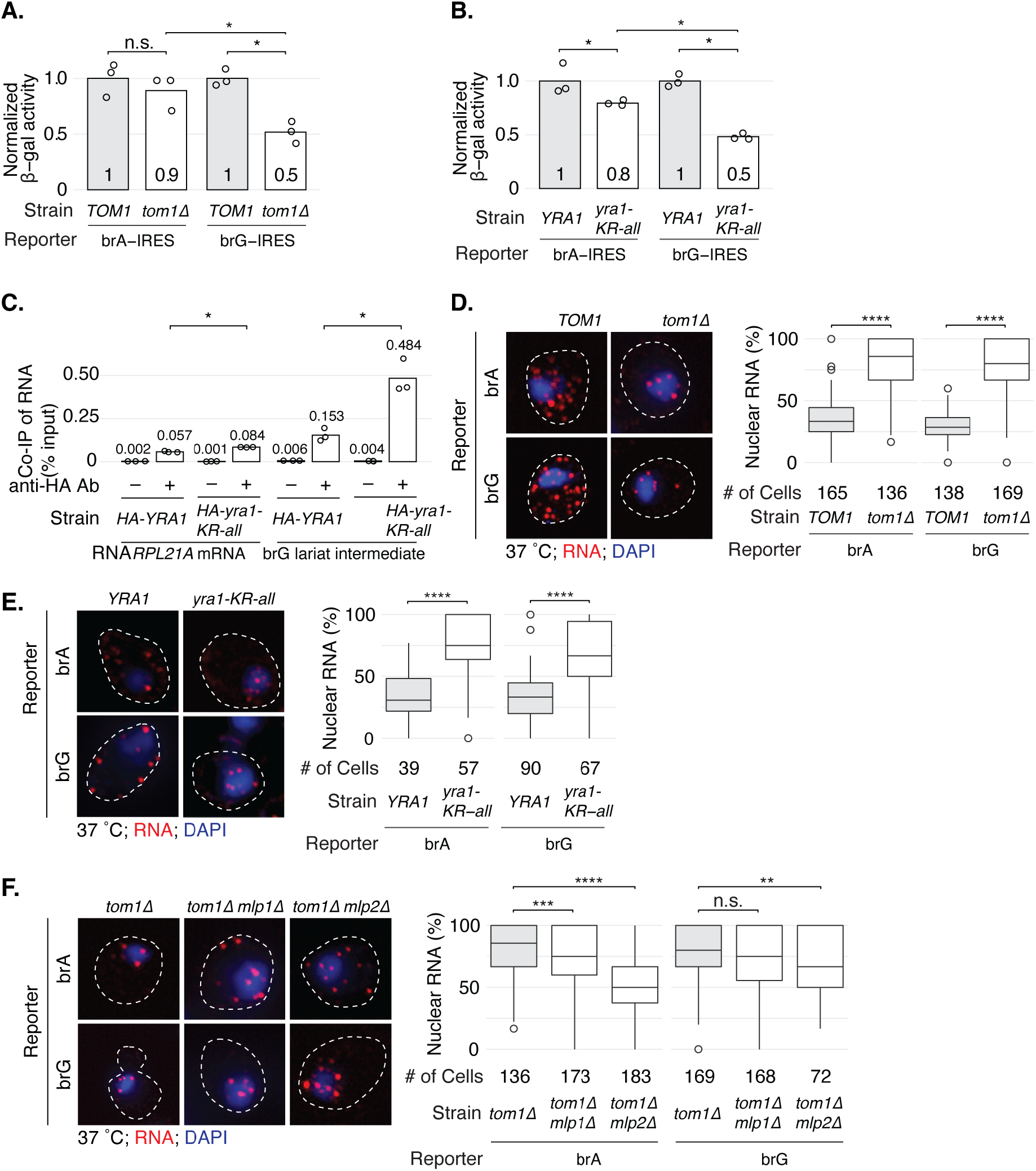
The export of lariat intermediate requires Tom1p-mediated ubiquitylation of Yra1p. **A, B.** The efficient export of lariat intermediates requires *TOM1* (**A**) and sites of *TOM1*-dependent ubiquitylation in *YRA1* (**B**). *TOM1* and *tom1Δ* cells (**A**) were grown at a permissive temperature of 30 °C, and *YRA1* and *yra1-KR-all* cells (**B**) were grown at semi-permissive temperature of 25 °C. The cytoplasmic localization of the reporters was assayed by measuring β-galactosidase activity of cell extracts. The activity of three technical replicates was quantitated and illustrated as in Fig. 3B; values for each reporter were normalized to the mean activity of the same reporter in the *TOM1* strain (**A**) or *YRA1* stain (**B**). **C.** The release of Yra1p from lariat intermediate, as for mRNA, requires sites of *TOM1*-dependent ubiquitylation. Crosslinking and RNA co-IP were executed as in Fig. 2C, except that both *YRA1-HA* and *yra1-KR-all-HA* cells were shifted to 37 °C for 2 hours before crosslinking; the levels of Immunoprecipitated RNA were quantitated as in Fig. 2C. **D, E.** Export of lariat intermediate, as well as mRNA, requires Tom1p (**D**) and ubiquitylation of Yra1p (**E**). Compromising the activities of these proteins accumulates lariat intermediate and mRNA in the nucleus. Cells were shifted to the non-permissive temperature of 37 °C for 2 hours and then assayed for reporter localization by smRNA FISH (left); the nuclear fraction of reporter RNA was quantitated (right). **F**. The nuclear basket factors *MLP1* and/or *MLP2* are required for the nuclear localization of mRNA and lariat intermediate. Cells were shifted to the non-permissive temperature of 37 °C for 2 hours and then assayed for reporter localization by smRNA FISH (left); the nuclear fraction of reporter RNA was quantitated (right). Throughout, for smRNA FISH cells were probed and demarked as in Fig. 1C; the nuclear fraction was calculated and displayed as in Fig. 1C.

Given this evidence that Tom1p is required specifically for the export of the brG-IRES reporter at the permissive temperature, we next tested whether ubiquitylation of the Tom1p target Yra1p is also required, exploiting the mutant *yra1-KR-all*, which mutates all potential ubiquitylation sites and precludes Tom1p-mediated ubiquitylation. At the semi-permissive temperature of 25 °C, the *yra1-KR-all* mutation reduced the β-galactosidase activity of the brA reporter but by a modest degree (20%; **Fig. 5B**), in contrast with the *mlp1Δ, nab2-F72AF73A*, and *tom1Δ* mutants (**Fig. 3**; **Fig. 4B,C**; **Fig. 5A**); this reduction may reflect the growth defect of this strain at 25 °C and/or the mutation of all lysines, which are not all functional targets for Tom1p (**Fig. S5**; Iglesias et al., 2010). The *yra1-KR-all* mutation reduced the β-galactosidase activity of the brG-IRES reporter by a greater degree (52%; **Fig. 5B**). Notably, the *mlp1Δ*, *nab2-F72A/F73A, tom1Δ*, and *yra1-KR-all* mutants all reduced the β-galactosidase activity of the brG-IRES reporter by a similar degree, roughly 50% (**Fig. 3B**; **Fig. 4B**; **Fig. 5A, B**), suggesting that Nab2p, Mlp1p, Tom1p, and Yra1p act in the same pathway.

Because Tom1p-mediated ubiquitylation of Yra1p promotes dissociation of Yra1p from its bound mRNPs before nuclear exit (Iglesias et al., 2010), we examined whether Yra1p ubiquitylation modulated the association of Yra1p with lariat intermediates, as documented in **Fig. 2C**, at 25 °C. By formaldehyde cross-linking followed by RNA co-IP using HA-tagged Yra1p, the *yra1-KR-all* mutation increased binding, as expected, to a control mRNA (*RPL21A*) by 1.5-fold; significantly, the *yra1-KR-all* mutation increased binding to the brG lariat intermediate by 3.2-fold (**Fig. 5C**). These data support a role for Tom1p-mediated ubiquitylation of Yra1p in promoting the export of lariat intermediates through dissociation of Yra1p.

Although Tom1p-mediated ubiquitylation of Yra1p has been found to promote its dissociation from mRNP before nuclear exit, it is still not clear whether Tom1p functions before or after mRNPs dock onto the nuclear pores. Genetic evidence has suggested that Tom1p acts after mRNPs dock to the nuclear pores (Iglesias et al., 2010). However, live tracking of newly born mRNAs from an induced reporter implied that Tom1p promotes mRNP docking, suggesting that Tom1p acts before mRNP docking (Saroufim et al., 2015). To distinguish these two models, we first examined directly the localization of lariat intermediates and mRNAs. By smRNA FISH, after a temperature shift of *tom1Δ* to the non-permissive temperature of 37 °C, the brG reporter, as well as the brA reporter, localized primarily to the nucleus (78% ±2% and 82% ±2%, respectively), relative to the wild-type strain (35% ±2% and 30% ±1%, respectively; **Fig. 5D**); similarly, after a temperature shift of the *yra1-KR-all* strain to 37 °C, the brG reporter, as well as the brA reporter, localized primarily to the nucleus (68% ±3% and 74% ±3%, respectively), relative to the wild-type strain (32% ±2% and 34% ±3%, respectively; **Fig. 5E**). These data indicate that at a non-permissive temperature of 37 °C, Tom1p-mediated ubiquitylation of Yra1p is required for the efficient export of not just mature poly(A)+ mRNA but also an immature RNA.

Together with the observation that *MLP1* and *MLP2* mutations suppress *TOM1* and *YRA1* mutants (Iglesias et al., 2010), our smRNA FISH results suggest that both brA-derived mRNAs and brG-derived lariat intermediates in *tom1Δ* and *yra1-KR-all* mutant cells are stalled on the nuclear periphery, likely at the nuclear basket (**Fig. 5D, E**). To test this idea, we examined whether deletion of the nuclear basket in the *tom1Δ* strain would rescue the export defect. Consistent with the previous genetic data that deletion of *MLP2* rescues the temperature sensitivity of the *tom1Δ* strain (Iglesias et al., 2010), deletion of *MLP2* in the *tom1Δ* strain substantially rescued the export defect of the brA reporter, reducing nuclear localization from 82% ±2% to 55% ±2% and increasing cytoplasmic localization by 2.5-fold (*p*-value=2.2×10^-16^; **Fig. 5F**); similarly, deletion of *MLP1* in the *tom1Δ* strain modestly but significantly reduced nuclear localization from 82% ±2% to 74% ±2% and increased cytoplasmic localization by 1.4-fold (*p*-value=8.6×10^-4^; **Fig. 5F**). Interestingly, deletion of *MLP2* but not *MLP1* in the *tom1Δ* strain also slightly but significantly rescued the export defect of the brG reporter, reducing nuclear localization from 78% ±2% to 68% ±3% and increasing cytoplasmic localization by 1.5-fold (*p*-value=0.003; **Fig. 5F**). The smaller increase of the brG reporter in the cytoplasm is consistent with an essential requirement of Mlp1p, if not also Mlp2p, in exporting lariat intermediates but not mRNAs. These results indicate that Mlp1p, and to a lesser degree Mlp2p, do not retain incompletely processed lariat intermediates in the nucleus but instead promote the export of such species. Indeed, together these data support a model in which Mlp1p and Mlp2p promote the export of lariat intermediates by facilitating docking at the nuclear basket and then Tom1p-mediated ubiquitylation of Yra1p promotes undocking, rather than docking, and thereby nuclear pore entry.

## Discussion

In this work, using a combination of ensemble and single molecule approaches, we discovered that the export of spliceosome-discarded lariat intermediates requires the general export receptor Mex67p (**Fig. 1**) and three of its adaptors, Yra1p, Nab2p, and Npl3p (**Fig. 2**). These data establish that lariat intermediates utilize the same export machinery as mRNAs. Unexpectedly, we found that the purported quality control factor Mlp1p did not retain lariat intermediates in the nucleus but instead promoted the export of lariat intermediates and did so through its interaction with the export adaptor Nab2p (**Fig. 3**; **Fig. 4**), implying a role for the nuclear basket in recruiting lariat intermediates to the NPC. Further, the export of lariat intermediates also relies on the E3 ubiquitin ligase Tom1p and its target sites in Yra1p (**Fig. 5**). Importantly, our findings imply that Tom1-mediated ubiquitylation of Yra1p undocks lariat intermediates from the nuclear basket of the NPC, allowing not only lariat intermediates but also mRNAs to transit through the NPC. Together, our results challenge a general role for Mlp1p in the quality control of mRNA export and implicate novel steps of mRNA export at the nuclear basket.

Our results establish that the general mRNA export pathway transports discarded lariat intermediates into the cytoplasm. Given that the mRNA export machinery assembles co-transcriptionally (Wende et al., 2019), it is likely that lariat intermediates are already export-competent when they are discarded by the spliceosome and thereby export by default. Our data further implicate the mRNA export pathway as a means for turning over RNA substrates discarded by the spliceosome. Indeed, a range of suboptimal pre-mRNAs and splicing intermediates are exported to the cytoplasm for degradation (Harigaya and Parker, 2012; Hilleren and Parker, 2003; Legrain and Rosbash, 1989; Mayas et al., 2010; Sayani and Chanfreau, 2012; Talhouarne and Gall, 2018); paralleling our own findings, the export of a fraction of pre-mRNA that accumulates upon pharmacological inhibition of the splicing factor SF3b1 requires the Yra1p human ortholog ALYREF (Carvalho et al., 2017). Such nuclear export may ensure that once substrates are discarded by the spliceosome, they are never engaged by the spliceosome again. Future work is needed to further understand the interplay between nuclear export and splicing proofreading.

Together with previous evidence of the export of unspliced pre-mRNAs and lariat intermediates (Carvalho et al., 2017; Harigaya and Parker, 2012; Hilleren and Parker, 2003; Legrain and Rosbash, 1989; Mayas et al., 2010; Sayani and Chanfreau, 2012; Talhouarne and Gall, 2018), our results presented here question a general role for Mlp1p in retaining immature mRNPs in the nucleus. Similarly, a genome-wide study assaying the export of pre-mRNA expressed from endogenous genes rather than reporters found no evidence that Mlp1p acts as a general retention factor (Sayani and Chanfreau, 2012). Further, recent studies indicate a positive role for TPR in mammals in promoting the export of a subset of mature mRNAs that are short, intron poor, or GC poor (Lee et al., 2020; Umlauf et al., 2013; Zuckerman et al., 2020). We note that in two studies providing some evidence for a role for TPR in nuclear retention, TPR knockdown increased total RNA levels of the reporter and, where analyzed, in both the cytoplasm and the nucleus (Coyle et al., 2011; Rajanala and Nandicoori, 2012), suggesting the possibility that TPR knockdown does not increase the export of nuclear RNAs but rather increases nuclear RNA levels and thereby indirectly leads to increased cytoplasmic levels. Indeed, TPR knockdown can impact nuclear and cytoplasmic RNA reporter levels without impacting overall, steady-state TPR levels and instead only affecting new TPR protein synthesis (Coyle et al., 2011), and even rapid depletion of TPR impacts the transcription of genes (Aksenova et al., 2020). By contrast, in our study, deletion of *MLP1* does not alter levels of pre-mRNA, lariat intermediate, or mRNA (**Fig. S2B, C; Fig. S3**), and we provide evidence that the impact of *MLP1* on lariat intermediate export reflects specifically an interaction between the C-terminus of Mlp1p and the Mlp1p-interacting domain of the mRNA export adapter Nab2p (**Fig. 4**), together providing support for a direct role for Mlp1p in exporting lariat intermediates. Given that a *pml39Δ* mutant phenocopies the retention defects of an *mlp1Δ* mutant and that Mlp1p is required for the localization of Pml39p to the nuclear basket (Palancade et al., 2005), it is formally possible that Pml39p instead of or in addition to Mlp1p plays a direct role in the export of lariat intermediates; however, Pml39p is also reciprocally required for the localization of Mlp1p to the nuclear basked, and Pml39p is not known to interact with Nab2p or with any other protein factor (Gunkel et al., 2023).

While both mRNAs and lariat intermediates require the general export pathway (Mex67p and its adapters; **Figs. 1 and 2**), only lariat intermediates seemingly require Mlp1p (**Fig. 3**). These results raise two possibilities: 1) Mlp1p functions specifically in the lariat intermediate export pathway, or 2) Mlp1p functions in both pathways but its role in mRNA export is masked by functional redundancy in the mRNA export pathway. The second possibility is more consistent with previous findings, which found Mlp1p and its orthologs promote mRNA export in a number of organisms (Aksenova et al., 2020; Bae et al., 2009; Lee et al., 2020; Li et al., 2021; Shibata et al., 2002; Umlauf et al., 2013; Xu et al., 2007; Zuckerman et al., 2020). Considering that multiple interactions form between an exporting mRNA and the NPC, we reason that, in *mlp1Δ* cells, other functionally redundant factors compensate for the loss of Mlp1p to ensure mRNA export. Therefore, the loss of one interaction, such as deletion of *MLP1*, is unlikely to grossly affect mRNA export in budding yeast. By contrast, the lariat intermediate does not have a 5’ cap, which facilitates mRNA export (Ashkenazy-Titelman et al., 2020), thus potentially rendering the export of this suboptimal export substrate more sensitive to disruptions, such as deletion of *MLP1*. Consistent with the idea that suboptimal export substrates may be more dependent on Mlp1p, the export of short, intron poor, or GC poor transcripts are dependent on TPR (Lee et al., 2020; Umlauf et al., 2013; Zuckerman et al., 2020). Suggesting a role for *MLP1/2* in docking mRNA at the nuclear basket, as well as a role for Tom1p in undocking mRNA (see below), we found that when *TOM1* is deleted mRNA is retained in the nucleus in an *MLP1-* and *MLP2*-dependent manner (**Fig. 5**; see below). Additionally, *MLP1* extends the residency time of mRNAs at the NPC (Saroufim et al., 2015). Further, TPR promotes engagement of mRNA with the nuclear basket (Li et al., 2021), potentially by recruiting mRNA via the TREX-2 component GANP (Aksenova et al., 2020). Lastly, the export of Balbiani ring RNPs in *Chironomus* begins with engagement of the ring of the nuclear basket (Wurtz-T et al., 1996), where the C-terminus of Mlp1p, the Nab2p-interacting domain in budding yeast, resides (Akey et al., 2022).

Notably, an exporting mRNA first docks to the nuclear basket of the NPC in an Mlp1p/TPR-dependent manner and then somehow transitions to engage the FG-repeat containing NUP153 deeper in the nuclear basket before entering the central channel of the NPC (Li et al., 2021; Saroufim et al., 2015). Our results suggest that Tom1p undocks exporting mRNAs from the Mlp1/2p and TPR components of the nuclear basket by ubiquitylating Yra1p. First, we and others have shown by RNA FISH that *tom1Δ* cells accumulate mRNA in the nucleus (**Fig. 5**) (Iglesias et al., 2010). Importantly, consistent with previous genetic evidence (Iglesias et al., 2010), we show directly that deletion of *MLP1* or especially *MLP2* rescues this export defect of mRNA in *tom1Δ* cells (**Fig. 5**). Given that Mlp1p and Mlp2p are subunits of the nuclear basket, these results imply that Tom1p acts at the nuclear basket, and that deletion of *MLP2* in particular renders Tom1p-mediated undocking unnecessary by bypassing mRNA docking to the nuclear basket. This view of a role for Mlp1/2p in docking and a role for Tom1p in undocking, provides an alternative to the view that Mlp1/2p and TPR perform quality control functions (Iglesias et al., 2010). As noted above, our data suggest that the nuclear basket can provide an essential docking role in export, as in the case of lariat intermediates, and that this role is redundant for other cargos, as in the case of our mRNA and pre-mRNA reporters; in the case of short, intron-poor, or GC-poor mRNAs, which require TPR for export (Lee et al., 2020; Umlauf et al., 2013; Zuckerman et al., 2020), such redundancy may be lacking. Most importantly, whether or not docking is essential, our data suggest that once cargo is docked, undocking is essential, rationalizing nuclear accumulation of both lariat intermediate and mRNA in *tom1Δ* or *yra1-KR-all* cells (**Fig. 5D, E**). This view would indicate that previous data suggesting the popular interpretation that the nuclear basket serves as a checkpoint (Coyle et al., 2011; Galy et al., 2004; Palancade et al., 2005; Rajanala and Nandicoori, 2012) instead imply that unspliced pre-mRNA at the nuclear basket is simply stuck on the mRNA export pathway and unable to undergo undocking (Bonnet and Palancade, 2015), as our evidence suggests for mRNA in the *tom1Δ* and *yra1-KR-all* mutants (**Fig. 5D, E**), and that deletion of *MLP1* or *MLP2* or depletion of TPR simply bypasses the requirement for undocking, as we observe for mRNA in the *mlp1Δ* or *mlp2Δ* mutants (**Fig. 5F**).

Collectively, our data suggest a model, in which lariat intermediates first dock onto the nuclear basket, requiring at least the interaction between Mlp1p and Nab2. Then, Tom1p is somehow activated to ubiquitylate Yra1 to undock lariat intermediates from the nuclear basket, allowing them to transit through the NPC, a pathway that likely also operates in mRNA export but in a manner that is normally masked by redundant pathways.

## Acknowledgements

We thank A. Corbett, F. Stutz, and T. Kress for sharing their reagents. We thank the members of the Staley lab for their helpful discussions, including Klaus Nielsen for comments on the manuscript. This work was funded by grant from the NIH (R01GM062264 to J.P.S.). All data, code, and materials used in the analysis are available upon request.

## Author contributions

Conceptualization, Y.Z. and J.P.S.; Experiments, data analysis, statistics, Y.Z.; interpretation, and visualization, Y.Z., and J.P.S.; funding acquisition, J.P.S.; writing – original draft, Y.Z. and J.P.S.; writing – review and editing, Y.Z. and J.P.S.

## Materials and Methods

### Yeast strain and plasmid construction

Yeast strains and plasmids used in this study are listed in table S1 and S2. Yeast strains were constructed with exogenous copy present on a shuffle plasmid. Plasmids were made either using topo cloning and/or site-directed mutagenesis. *pRS313-MLP1* plasmid was generated by cloning *MLP1* locus (ORF with 500 bp flanking sequences) from the *BY4741* strain into *pRS313* empty vector. *pRS313-mlp1-Δ1586-1768* plasmid was generated by site directed mutagenesis of *pRS313-MLP1*. The NLS in *pRS313-mlp1-Δ1586-1768* is intact. GFP tagged *MLP1* plasmids (*pRS313-MLP1-GFP* and *pRS313-mlp1-Δ1586-1768-GFP*) were generated by inserting the enhanced monomeric GFP sequence immediately downstream of the penultimate amino acid in frame. *pRS315-nab2-F72AF73A* plasmid was generated by site-directed mutagenesis of *pRS315-NAB2* plasmid. *pRS316-MEX67* plasmid was generated by cloning *MEX67* locus (ORF with 500 bp flanking sequences) from the *BY4741* strain into *pRS316* empty vector. *pRS313-MEX67* was generated by subcloning *MEX67* from *pRS316-MEX67* to *pRS313*. *mex67-5* mutant was generated by site directed mutagenesis of *pRS313-MEX67*. Reporters used in FISH experiments were generated as followed. The *lacZ* reporters containing different mutations were subcloned from *pRS426* to *pRS316*. Then, the strong *GPD* promoter was replaced by a weak *STE5* promoter. *MEX67* strains were generated as followed. *MEX67* shuffle strain was generated by replacing the endogenous *MEX67* with *HPHMX4* in the *BY4741* strain expressing *pRS316-MEX67*. *MEX67* shuffle strain was conferred by PCR and 5-FOA selection. Afterward, *pRS316-MEX67* in the *MEX67* shuffle strain was replaced by either *pRS313-MEX67* or *pRS313-mex67-5*, generating the final *MEX67* strain or *mex67-5* strain. Similar strategy was applied to generate *mlp1Δ dbr1Δ* strain from *dbr1Δ* strain, *mlp1Δ tom1Δ* strain from *tom1Δ*, and *mlp2Δ tom2Δ* strain from *tom1Δ* strain.

### RNA-FISH

RNA-FISH was performed as described in Raj and Tyagi, 2010 (Raj and Tyagi, 2010). Cells were imaged using Olympus IX81 inverted widefield microscope equipped with Hamamatsu Orca Flash 4.0 camera with 4 megapixels and 100x 1.45NA oil objective lens. Single RNA molecule counting was conducted using custom macros in imageJ and statistical analysis was conducted in R. 10% formamide was used for smRNA FISH experiments and 40% formamide was used for RNA-FISH experiments. For smRNA FISH, probes were designed by Biosearch targeting the *lacZ* portion of the reporters. For RNA-FISH, a single Cy3-labeled oligo-dT(50) probe was used to target polyA+ RNAs and a single Alexa488-labeled probe was used to target the brG reporter.

### Immunofluorescence

Cells expressing *MLP1-GFP* or *MLP1-Δ1586-1768-GFP* were imaged in an anti-fade buffer using epifluorescence microscopy with 100X magnification.

### RNA co-immunoprecipitation

For Mex67-GFP RNA co-IP, cell lysates were prepared from *MEX67-GFP* cells expressing the brG reporter (pJPS1488) and incubated with beads pre-conjugated with either anti-GFP antibody or IgG. After a 2-hour incubation at 4 °C, beads were washed five times and put through RNA extractions using phenol:chloroform:isoamyl alcohol (25:24:1, v/v). To ensure Mex67 binds lariat intermediates *in vivo*, tagged *MEX67-TAP* cells expressing the empty vector were mixed with untagged *MEX67* cells expressing the brG reporter. In parallel, tagged *MEX67-TAP* cells expressing the brG reporter were mixed with untagged *MEX67* cells expressing the empty vector. Lysates were then prepared from these mixed cells and incubated with IgG beads at 4 °C for 2 hours. After incubation, beads were washed five times. Bound *MEX67* containing mRNPs were released by TEV cleavage and put through RNA extractions using phenol:chloroform:isoamyl alcohol (25:24:1, v/v). For Nab2-HTB denaturing RNA co-IP and HA-Yra1 denaturing RNA co-IP, cells were grown to OD600 of 0.6-0.8, cross-linked for 10 minutes by 3.7% formaldehyde, and quenched with glycine. Then, cell lysates were prepared using glass beads. Cell lysates were incubated with either Ni-NTA beads in the case of Nab2-HTB or anti-HA beads in the case HA-Yra1 at 4 °C for 2 hours. After incubation, beads were washed five times and put through RNA extractions using phenol:chloroform:isoamyl alcohol (25:24:1, v/v). Lariat intermediates were assayed using lariat specific RT-PCR. mRNA of endogenous *RPL21* was assay by RT-PCR as an internal control.

### In vivo RNA analysis

For *in vivo* RNA analysis, cells transformed with ACT1-IRES-lacZ splicing reporters were cultured in selective media at 30 °C to an OD600 of 0.6–0.8, lysed, and then assayed for RNA. RNA was analyzed by primer extension using 32P-radiolabeled primers and AMV-reverse transcriptase, followed by the separation of products on a 6% denaturing polyacrylamide gel. Two primmer were used, one binding the *lacZ* portion of the reporters and the other binding U14 snoRNA for internal control. Data were visualized using a phosphorimager (Molecular Dynamics) and quantitated using ImageQuant (GE Healthcare).

### β-galactosidase assays

Liquid assays were performed as described in Mayas et al., 2010. Specifically, 1.5 mL of liquid cultures in selective media were harvested at OD 600 of 0.6–0.8, and washed with Z buffer (60 mM Na_2_HPO_4_, 40 mM NaH_2_PO_4_, 10 mM KCl, 1 mM MgSO_4_; pH 7.0). Cells were resuspended in 100 μL of Z buffer and lysed by 6 cycles of freeze-thawing (30 seconds each in liquid nitrogen and in a 42 °C water bath). Lysed cells were incubated with 700 μL of prewarmed (30 °C) Z buffer that included 1 mg mL *ortho*-nitrophenyl-β−galactopyranoside and 50 mM β-mercaptoethanol. Reactions were incubated at 30 °C for 30 minutes to 2 hours and stopped by adding 0.5 mL 1M Na_2_CO_3_. After cell debris was pelleted, the OD_420nm_ of the supernatant was measured. Activity in Miller units was calculated as (OD4200nm × 1000)/(OD600nm × (minutes elapsed) × 1.5 mL).

## Supplementary figures and tables

**Figure S1.**
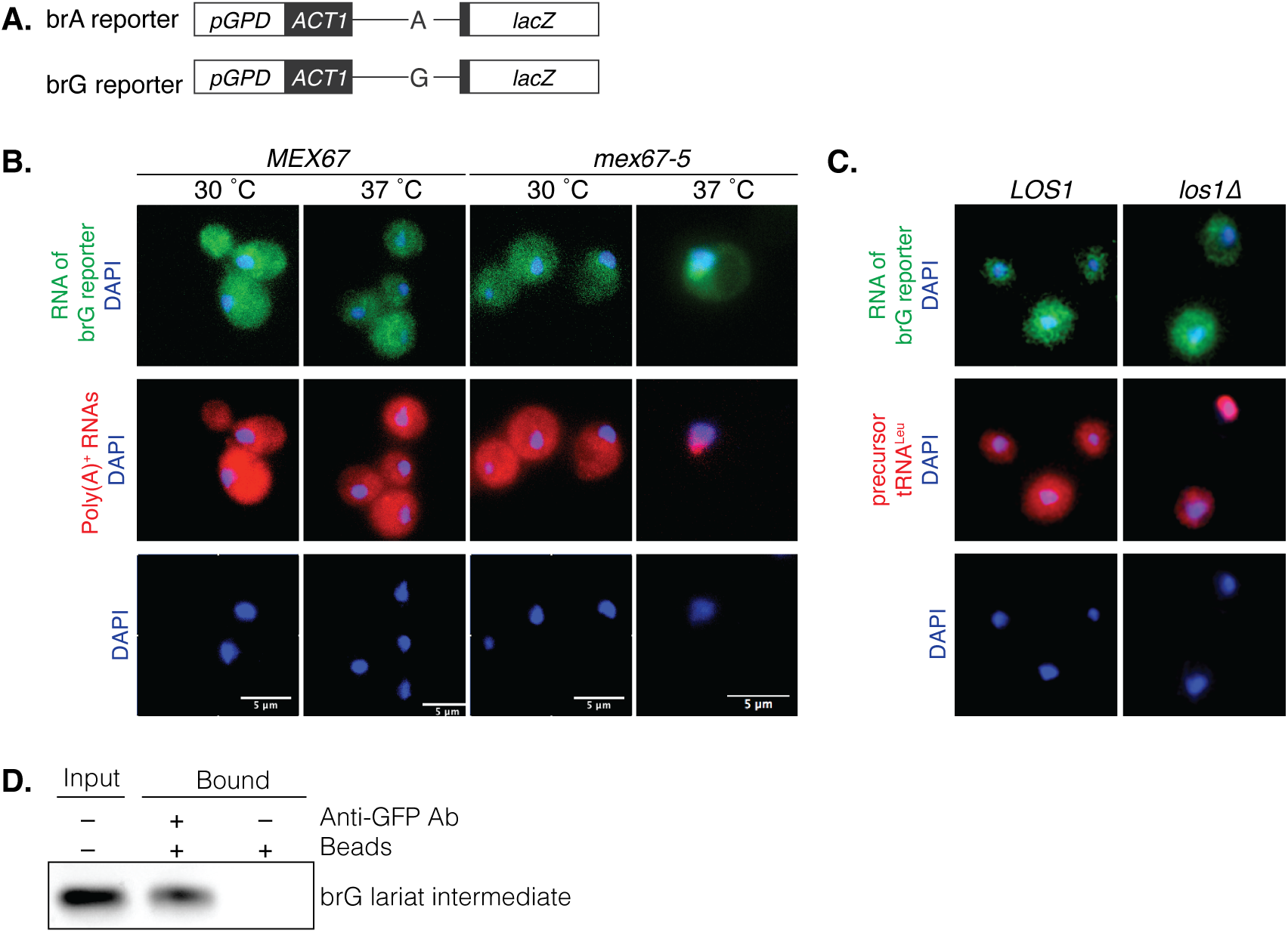
Export of a lariat intermediate requires the mRNA export factor Mex67p but not the tRNA export factor Los1; related to Fig. 1. **A.** Schematic representation of the brA and brG export reporters driven by *pGPD* promoter. **B.** By RNA-FISH, the export of the brG reporter, like poly(A)+ RNA, is impeded by the *mex67-5* mutant at the non-permissive temperature. Cells were shifted from 30 °C to 37 °C for 30 minutes. The intronic region of the brG reporter transcript was detected by Alexa Fluor 488-labelled probes; poly(A)+ RNA was detected in the same cells by a Cy3-labelled poly(dT)_50_ probe; DNA was probed by DAPI, which is shown separately and overlayed. **C.** By RNA-FISH, the export of the brG reporter, unlike pre-tRNA, is not impeded by the *los1Δ* mutant. The intronic region of the brG reporter was detected as in panel A; the intronic region of the tRNA^Leu^ precursor was detected in the same cells by a Cy3-labelled probe (note that pre-tRNA splicing does not involve the spliceosome and occurs in the cytoplasm in budding yeast); DNA was probed by DAPI, which is shown separately and overlayed. **D**. By native RNA co-IP, Mex67-GFP interacts with brG lariat intermediates, which were detected by RT-PCR using lariat specific primers.

**Figure S2.**
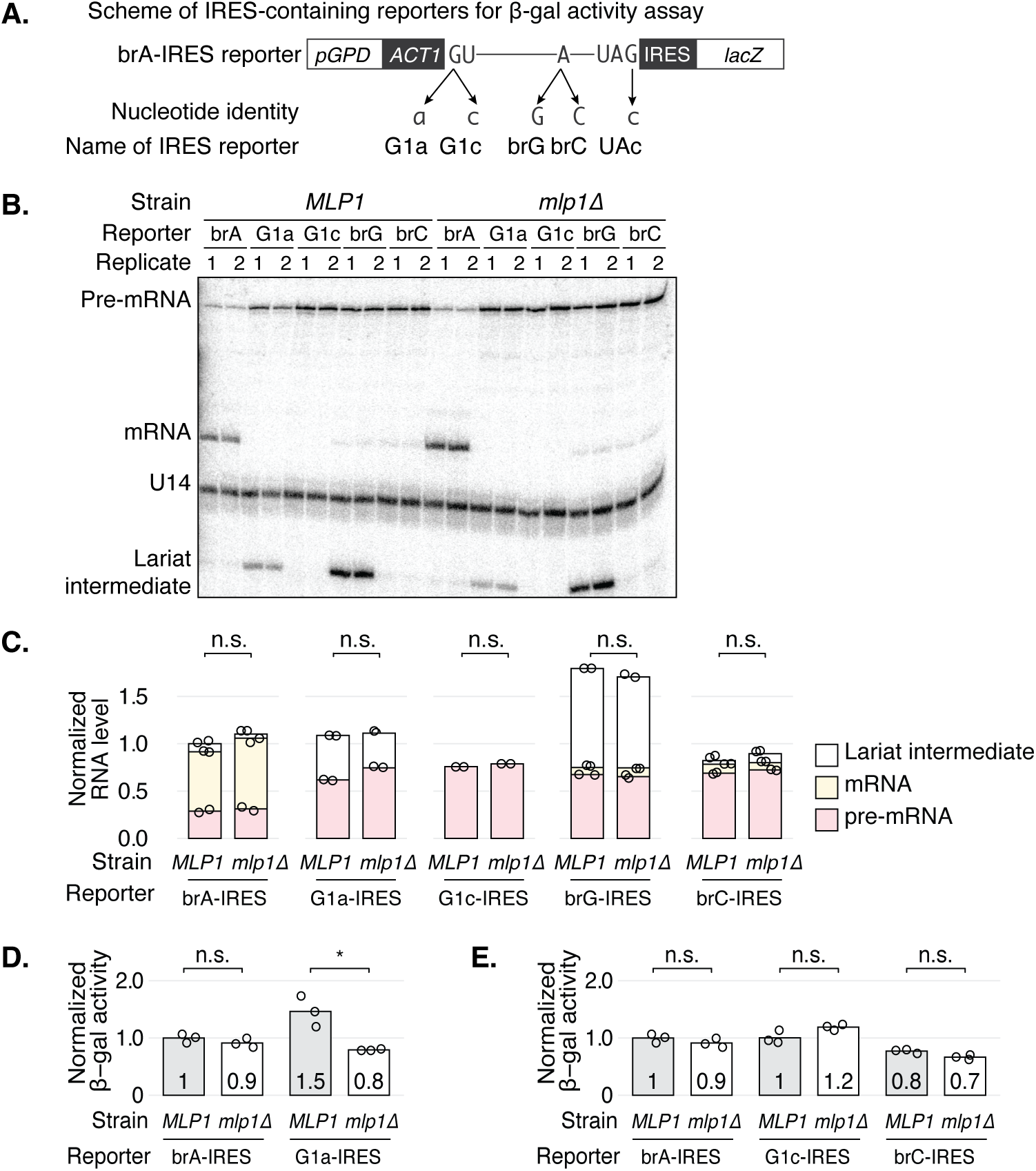
Whereas the export of lariat intermediates requires *MLP1*, the export of pre-mRNA does not; related to Fig. 3. **A**. Schematic representation of different *ACT1-IRES-lacZ* export reporters driven by the *pGPD* promoter. **B, C**. The *mlp1Δ* mutation does not significantly impact the levels of splicing species for the wild-type IRES-containing reporter (brA) or mutated reporters (brG, brC, G1a or G1c), ruling out a role for *MLP1* in splicing or RNA stability and consistent with a role for *MLP1* in the export of brG lariat intermediates. In panel **B**, the splicing of each reporter was analyzed in *MLP1* and *mlp1Δ* cells, grown at 30 °C. The levels of splicing precursor, lariat intermediate, and mRNA, were detected by extension of a primer that annealed to the downstream exon; the levels of the snoRNA U14 were detected similarly and served as an internal control; two technical replicates are shown. Panel **C** shows normalized RNA levels of different splicing species from the two technical replicates in B; the bar height indicates the mean RNA level. Levels of pre-mRNA (salmon), lariat intermediate (yellow), and mRNA (white) were stacked to reflect total 3′ exon levels and normalized to RNA levels of the *brA-IRES-lacZ* reporter in the *MLP1* strain. **D**. The export of the G1a lariat intermediate is comprised by the *mlp1Δ* mutation. *MLP1* or *mlp1Δ* cells were grown at 30 °C, and the cytoplasmic localization of the *brA-* and *G1a-IRES-lacZ* reporters was assayed by measuring β-galactosidase activity of cell extracts. The activity was quantitated as in Fig. 3B; values were normalized to the *brA-IRES-lacZ* reporter in the *MLP1* strain. **E**. The export of pre-mRNA is not comprised by the *mlp1Δ* mutation. *MLP1* or *mlp1Δ* cells were grown at 30 °C, and the cytoplasmic localization of the *brA-, G1c-* and *brC-IRES-lacZ* reporters was assayed by measuring β-galactosidase activity of cell extracts. The activity was quantitated as in Fig. 3B; values were normalized to the brA-IRES reporter in the *MLP1* strain. The activities for brA reporter were reproduced from **D**.

**Figure S3.**
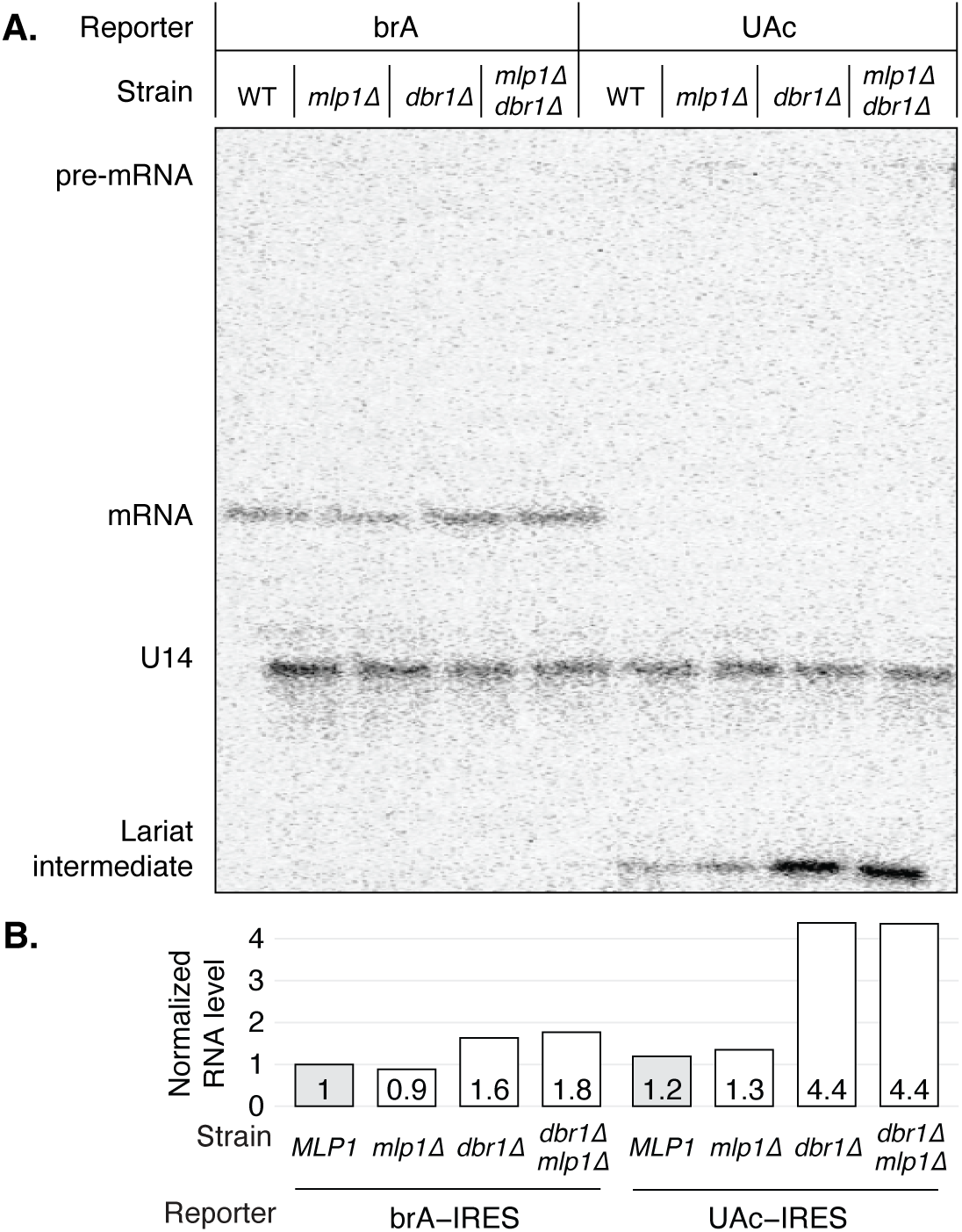
The *mlp1Δ* mutation does not impact the levels of splicing species for either the brA- or UAc-IRES reporters in the presence or absence of Dbr1p; related to Figure 3. In panel **A**, *MLP1*, *mlp1Δ*, *dbr1Δ*, or *mlp1Δ dbr1Δ* cells were grown at 30 °C and splicing of the reporters was analyzed by primer extension, as described in panel Fig. S2B. Panel **B** shows the quantitation of total RNA levels. Values were normalized to the brA-IRES reporter in the *MLP1* strain. Note that the lariat intermediate dominates the splicing species from the UAc-IRES reporter in the *dbr1Δ* strain, where this species is specifically stabilized.

**Figure S4.**
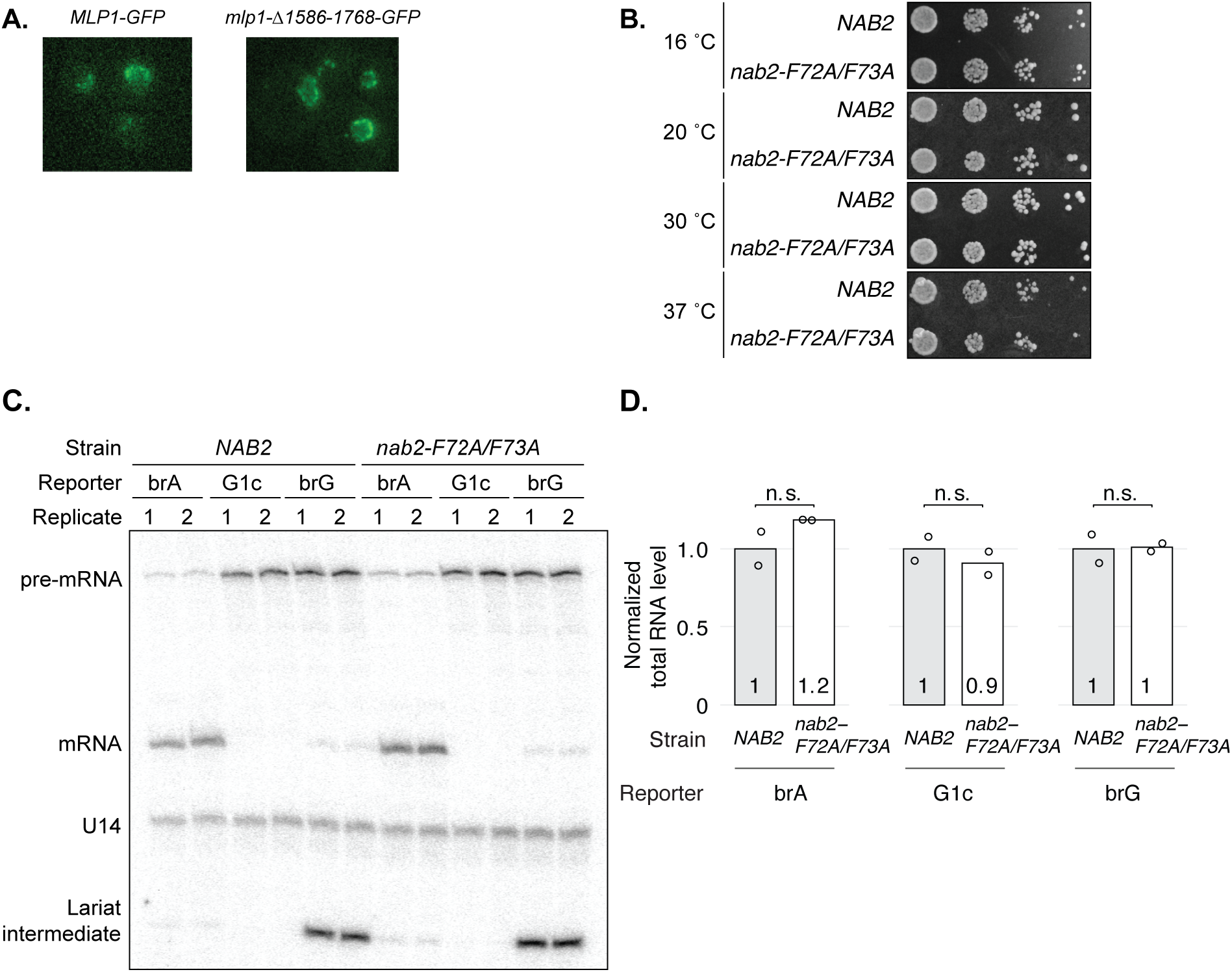
Deletion of Nab2-interacting domain in Mlp1p does not compromise Mlp1p localization, and nab2-F72A/F73A does not compromise either growth or splicing; related to Figure 4. **A.** Mutated mlp1p-Δ1586-1768-GFP localizes to the nuclear periphery, just as wild-type Mlp1p-GFP does. GFP was probed (in live cells) by epi-fluorescence microscopy. Note that the crescent shape is characteristic of Mlp1p localization, which reflects exclusion from the region of the nuclear periphery where the nucleolus resides. Also note that whereas mlp1p-Δ1586-1768-GFP lacks the Nab2p-interacting domain, the mutated protein does include its nuclear localization signal. **B.** The *nab2-F72A/F73A* mutant displays no growth defect over a range of temperatures. Growth of the indicated strains was assayed at the indicated temperatures by spotting serial dilutions of cultures onto rich, solid media (YPDA plates) and incubating for 5 days at 16 °C, 4 days at 20 °C, 2 days at 30 °C, and 1 day at 37 °C. **C, D.** The *nab2-F72A/F73A* double mutation does not impact the splicing of the brA, G1c, or brG reporters. In panel **C**, splicing was analyzed by primer extension as in Fig. S2B. Panel **D** shows the quantitation of total RNA levels. The total RNA of each reporter was normalized to its RNA level in the *NAB2* strain.

**Figure S5.**
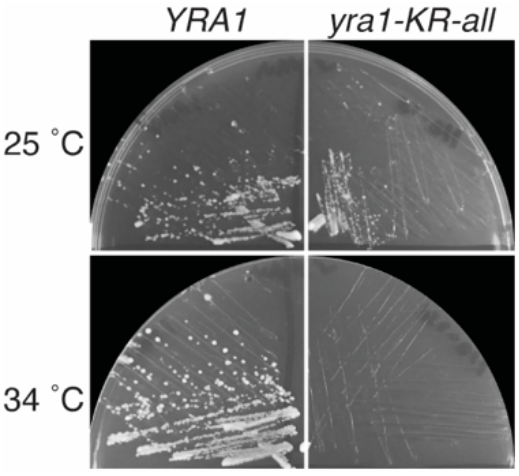
The yra1-KR-all mutant displayed a mild growth defect at 25 °C; related to Figure 5. *YRA1* and *yra1-KR-all* cells were streaked onto YPDA, solid media, and incubated for 2 days at 25 °C or 34 °C.

**Table S1.**
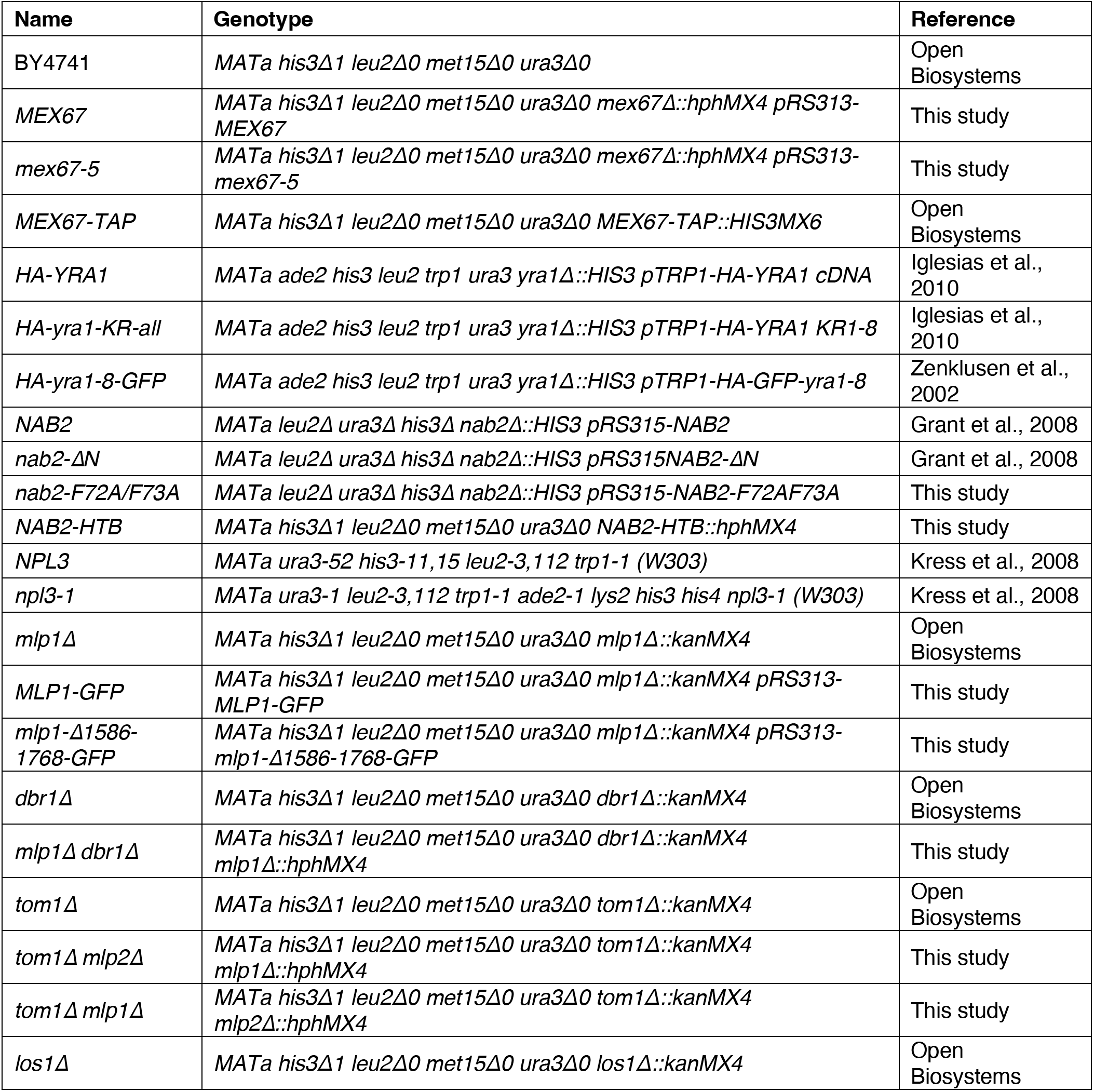
Yeast strains used in this study.

**Table S2.** Plasmids used in this study.

| Plasmid ID | Name | Reference |
| --- | --- | --- |
| pYZ1 | *pRS313-MLP1* | This study |
| pYZ4 | *pRS313-mlp1-Δ1586-1768* | This study |
| pYZ9 | *pRS313-MLP1-GFP* | This study |
| pYZ10 | *pRS313-mlp1-Δ1586-1768-GFP* | This study |
| pAC717 | *pRS315-NAB2 (pAC717)* | Green et al., 2002 |
| pYZ25 | *pRS315-nab2-F72AF73A (derived from pAC717)* | This study |
| pYZ17 | *pRS315-MEX67* | This study |
| pYZ18 | *pRS315-mex67-5* | This study |
| pYZ19 | *pRS316-MEX67* | This study |
|  | *pUN100-LEU2-MEX67-GFP* | Segref et al., 1997 |
| pJPS1481 | *pRS426-GPDpromoter-ACT1(WT)-IRES-lacZ* | Mayas et al., 2010 |
| pJPS1483 | *pRS426-GPDpromoter-ACT1(G303c)-IRES-lacZ* | Mayas et al., 2010 |
| pJPS1485 | *pRS426-GPDpromoter-ACT1(G1a)-IRES-lacZ* | Mayas et al., 2010 |
| pJPS1486 | *pRS426-GPDpromoter-ACT1(G1c)-IRES-lacZ* | Mayas et al., 2010 |
| pJPS1488 | *pRS426-GPDpromoter-ACT1(brG)-IRES-lacZ* | Mayas et al., 2010 |
| pJPS1514 | *pRS426-GPDpromoter-ACT1(brC)-IRES-lacZ* | Mayas et al., 2010 |
|  | *pRS316-STE5promoter-ACT1(WT)-IRES-lacZ* | This study |
|  | *pRS316-STE5promoter-ACT1(brG)-IRES-lacZ* | This study |
|  | *pRS316-STE5promoter-ACT1(G303c)-IRES-lacZ* | This study |
|  | *pRS316-STE5promoter-ACT1(WT)-lacZ* | This study |
|  | *pRS316-STE5promoter-ACT1(brG)-lacZ* | This study |

